# A multiscale closed-loop neurotoxicity model of Alzheimer’s disease progression explains functional connectivity alterations

**DOI:** 10.1101/2023.09.24.559180

**Authors:** Jesús Cabrera-Álvarez, Leon Stefanovski, Leon Martin, Gianluca Susi, Fernando Maestú, Petra Ritter

**Affiliations:** Complutense University of Madrid: Universidad Complutense de Madrid; Charité- Universitätsmedizin Berlin

## Abstract

While the accumulation of amyloid-beta (*Aβ*) and hyperphosphorylated-tau (hp-tau) as two classical histopathological biomarkers are crucial in Alzheimer’s disease (AD), their detailed interaction with the electrophysiological changes at the meso- and macroscale are not yet fully understood. We developed a mechanistic mequltiscale model of AD progression, linking proteinopathy to its effects on neural activity and vice-versa. We integrated a heterodimer model of prion-like protein propagation, and a network of Jansen-Rit electrical oscillators whose model parameters varied due to neurotoxicity. Changes in inhibition guided the electrophysiological alterations found in AD, and both *Aβ* and hp-tau-related inhibition changes were able to produce similar effects independently. Additionally, we found a causal disconnection between cellular hyperactivity and interregional hypersynchrony. Finally, we demonstrated that early *Aβ* and hp-tau depositions’ location determine the spatiotemporal profile of the proteinopathy. The presented model combines the molecular effects of both*Aβ* and hp-tau together with a mechanistic protein propagation model and network effects within a unique closed-loop model. This holds the potential to enlighten the interplay between AD mechanisms on various scales, aiming to develop and test novel hypotheses on the contribution of different AD-related variables to the disease evolution.

**Significance Statement:** This research presents a groundbreaking closed-loop model of AD mechanisms, bridging the gap between protein distribution and neural activity. Contrary to prior assumptions, the study reveals that interregional hyper-synchrony and cellular hyperactivity are not directly linked. Notably, the model identifies neural inhibition as a potential causal factor in neurophysiological AD alterations and posits early depositions of *Aβ* as a determinant of the spatiotemporal profile of proteinopathy. The significance of this mechanistic disease framework lies in its potential to produce insights into AD evolution and to guide novel treatment strategies. It underscores the importance of further experiments and modelling efforts to refine our understanding of AD, offering hope for more effective treatments and personalized care in the fight against dementia.

## 1 Introduction

Alzheimer’s disease (AD) is a progressive neurodegenerative disorder widely characterized at several levels of analysis from molecules to cells to the whole brain. The interactions between these levels of analysis remain poorly understood, contributing to the limited treatment options (Breijyeh and Karaman, 2020; van Dyck et al., 2023). A mechanistic understanding of AD is essential for developing novel therapeutic interventions.

The main pathophysiological changes in AD involve the misfolding and accumulation of amyloid-*β* (*Aβ*) and a hyperphosphorylated version of the tau protein (hp-tau) in the brain (Masters et al., 2015). These two proteins have different toxic effects, while *Aβ* generates hyperactivity through the disruption of GABAergic inhibitory synapses and the reduction of glutamate reuptake, hp-tau disrupts the synaptic connectivity of neurons by reducing the number of dendritic spines in pyramidal cells Maestú et al. (2021). *Aβ* tends to aggregate into plaques, while hp-tau aggregates into neurofibrillary tangles (NFTs). Following the prion hypothesis of AD, misfolded *Aβ* and hp-tau propagate through the brain acting as seeds (i.e., prions) that trigger the misfolding and aggregation of their normal counterparts (Frost and Diamond, 2009; Gomez-Gutierrez and Morales, 2020; Walker, 2018). It has been shown that the hyperactivity produced by *Aβ* orients the prionic propagation of hp-tau in the brain (Rodriguez et al., 2020) following the Braak stages from the entorhinal cortex to the neocortex, producing a network disruption that has been linked to cognitive decline (Braak and Braak, 1991; Braak et al., 2011; Therriault et al., 2022). The higher the activity levels in a region, the faster the propagation of hp-tau to those regions. Additionally, hyperactivation has also been linked to an enhanced extracellular secretion and deposition of *Aβ* generating a cyclical phenomenon that reinforces bidirectionally neural hyperactivity and *Aβ* concentration (Cirrito et al., 2005; Yamamoto et al., 2015; Kamenetz et al., 2003; Busche and Hyman, 2020; Stargardt et al., 2015).

At the whole brain level, electrophysiological recordings have shown an early increase in alpha power and hypersynchrony between parietal regions (Nakamura et al., 2017, 2018), followed by a disruption in those network’s functional connectivity (FC) and a rise in anterior hypersynchrony (López-Sanz et al., 2016; Pusil et al., 2019). Finally, the whole network gets disrupted due to the effect of hp-tau accumulation and further generation of NFTs (Pereira et al., 2019; Ranasinghe et al., 2014; Ahnaou et al., 2017; Palop et al., 2006). Whether the hypersynchronization is linked to the previously mentioned cellular hyperactivity remains unknown (Maestú et al., 2021). Here, we hypothesize that hyperactivity causes an increase in inter-regional synchrony (i.e., FC) due to a higher rate of neural communication that enhances the coordination of distant neuronal groups.

In recent years, significant computational modelling efforts have employed brain network models (BNMs) to investigate neural activity and its relationship to several aspects of the disease (Stefanovski et al., 2021, 2019; Alexandersen et al., 2023; van Nifterick et al., 2022; Ranasinghe et al., 2022). These models reproduce neural activity by using sets of second-order differential equations known as neural mass models (NMMs) that are interconnected through empirically-derived structural connectivity (SC) networks based on tractography. These models are expected to serve as a tool for getting deeper insights into the mechanisms of AD and to be able to predict its appearance and evolution.

In this study, we constructed a multiscale neurotoxicity model based on neuroimaging data that includes molecular, cellular, and interregional features of the brain. Specifically, it integrates a heterodimer model for the generation and propagation of AD-related proteins, neuronal population activity, and interregional synaptic coupling. We aimed to explore the mechanisms that give rise to the observed neurophysiological changes in AD and to evaluate the relative impact of the different proteins involved. To pursue these goals, first, we explored an isolated neural population and the effects produced by changing the parameter candidates affected by AD (i.e., excitation and inhibition-related parameters). Second, we simulated the neurotoxicity model and adjusted its parameters to reproduce the empirically observed phenomena in AD (i.e., frequency slowing, relative alpha decrease, increase/decrease in excitation, and increase/decrease in functional connectivity). Finally, we use the resulting network model to analyze the impact of different biological mechanisms giving rise to AD progression. The mechanistic explanations derived from the model are key to understanding the effects of delivered treatments, fostering the development of new ones, and enhancing early detection methods.

## 2 Methods

### 2.1 Data acquisition

Magnetic resonance imaging (MRI) scans were acquired for 20 healthy participants (17 females) with mean age 60.5 (sd 4.17) at the [Author University Redacted]. Diffusion-weighted images (dw-MRI) were acquired with a single-shot echo-planar imaging sequence with the parameters: echo time/repetition time = 96.1/12,000 ms; NEX 3 for increasing the SNR; slice thickness = 2.4 mm, 128×128 matrix, and field of view = 30.7 cm yielding an isotropic voxel of 2.4 mm; 1 image with no diffusion sensitization (i.e., T2-weighted b0 images) and 25 dw-MRI (b = 900 s/mm2).

All participants provided informed consent.

### 2.2 Structural connectivity

In a preprocessing stage, FSL eddy was used to correct for eddy current distortion. The correction was conducted through the integrated interface in DSI Studio (http://dsi-studio.labsolver.org). Then, the eddy-corrected dw-MRI data was rotated to align with the AC-PC line. The restricted diffusion was quantified using restricted diffusion imaging (Yeh et al., 2016). The diffusion data were reconstructed using generalized q-sampling imaging (Yeh et al., 2010) with a diffusion sampling length ratio of 1.25. The tensor metrics were calculated using dw-MRI with a b-value lower than 1750 s/mm ^2^. A deterministic fibre tracking algorithm (Yeh et al., 2013) was used with augmented tracking strategies (Yeh, 2020) to improve reproducibility. The whole brain was used as seeding region. The anisotropy threshold was randomly selected. The angular threshold was randomly selected from 15 degrees to 90 degrees. The step size was randomly selected from 0.5 voxels to 1.5 voxels. Tracks with lengths shorter than 15 or longer than 300 mm were discarded. A total of 5 million seeds were placed.

A combination of Desikan-Killiany (Desikan et al., 2006) and ASEG (Fischl et al., 2002) atlases were used as a volume parcellation atlas, and two SC matrices were calculated: weights by using the count of the connecting tracks, and tract length by using the average length of the streamlines connecting two regions. The final SC consisted of a subset of these networks including 40 regions (see Table 3) representing the cingulum bundle (Bubb et al., 2018), one of the most prominent white matter structures that interconnect frontal, parietal, temporal and subcortical regions, and whose impairment has been related to AD and cognitive decline (Yu et al., 2017; Chen et al., 2023; Bubb et al., 2018).

### 2.3 Brain network model

SC matrices served as the skeleton for the BNMs implemented in TVB (Sanz-Leon et al., 2015) where regional signals were simulated using Jansen-Rit (JR) NMMs (Jansen and Rit, 1995). The JR is a biologically inspired model of a cortical column capable of reproducing alpha oscillations through a system of second-order coupled differential equations (see Table 1 for a description of parameters):

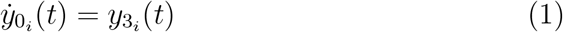

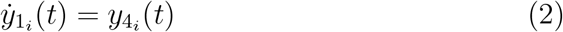

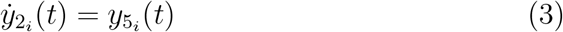

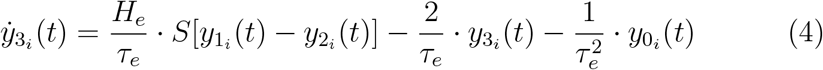

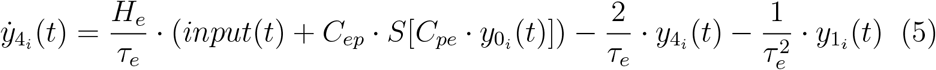

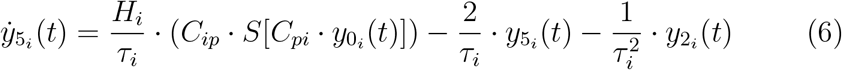

Where:

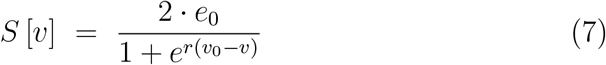

The inter-regional communication introduces heterogeneity in terms of connection strength *w*_*ji*_, and conduction delays *d*_*ji*_ between nodes i and j where: 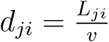, with *L*_*ji*_ being the length of the tract from node i to node j, and *v* representing the conduction speed. The input is given by:

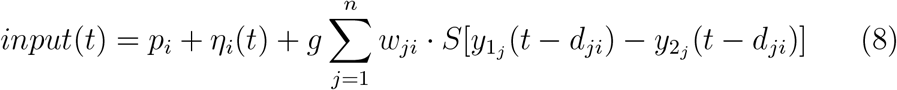

It represents the electrophysiological activity (in voltage) of three sub-populations of neurons: pyramidal neurons (*y*_0_), excitatory interneurons (*y*_1_), and inhibitory interneurons (*y*_2_) and their second derivatives (*y*_3_, *y*_4_, and *y*_5_, respectively). These subpopulations are interconnected and integrate external inputs from other cortical columns. The communication is implemented in terms of firing rate, and a sigmoidal function (eq. 7) stands for the conversion from voltage to firing rate.

**Table 1.** JR-BNM parameters used in simulations.

| Parameter | Value | Unit | Description |
| --- | --- | --- | --- |
| $H_e$ | 3.25 | mV | Average excitatory synaptic gain |
| $H_i$ | 22 | mV | Average inhibitory synaptic gain |
| $\tau_e$ | 10 | ms | Time Constant of excitatory PSP |
| $\tau_i$ | 20 | ms | Time Constant of inhibitory PSP |
| $C_{pe}$ | 135 | | Average synaptic contacts: pyramidals to excitatory interneurons |
| $C_{ep}$ | 108 | | Average synaptic contacts: excitatory interneurons to pyramidals |
| $C_{pi}$ | 33.75 | | Average synaptic contacts: pyramidals to inhibitory interneurons |
| $C_{ip}$ | 33.75 | | Average synaptic contacts: inhibitory interneurons to pyramidals |
| $e_0$ | 0.0025 | $ms^{-1}$ | Half the maximum firing rate |
| $r$ | 0.56 | $mV^{-1}$ | Slope of the presynaptic function at $v_0$ |
| $v_0$ | 6 | mV | Potential when half the maximum firing rate is achieved |
| $p$ | 0.1085 | $ms^{-1}$ | Mean of random gaussian intrinsic input |
| $\eta$ | 0.022 | $ms^{-1}$ | Standard deviation of random gaussian intrinsic input (noise) |
| $g$ | 25 | | Coupling factor for inter-regional communication |
| $s$ | 20 | mm/ms | Conduction speed for inter-regional communication |

The input represents two main drivers of activity in the NMMs: inter-regional communication and intrinsic input. The latter is defined by a Gaussian noise with *p* mean and *η* std.

A global coupling factor *g* is implemented to scale linearly tracts’ weights. Both, *g* and *v* are scaling factors that apply to all nodes and were adjusted to reproduce the qualitative features of the AD evolution (see Figure 4-1).

### 2.4 Closed-loop neurotoxicity model

We have adapted an already published model for AD protein propagation, first shown by Thompson et al. (2020) that reproduces the evolution of tau and *Aβ* concentrations in the AD brain, and a further extension by Alexandersen et al. (2023) that linked this evolution to changes in neural activity patterns using NMMs. Here, we adapt this latter model by implementing a JR NMM and introducing a feedback mechanism that connects the NMMs’ activity to the production and diffusion of toxic proteins in the AD brain.

**Table 2.** Parameters used in the closed-loop neurotoxicity model.

| Parameter | Value | Unit | Description |
| --- | --- | --- | --- |
| $\text{prod}_{A\beta}$ | 3 | $\text{M} \cdot \text{years}^{-1}$ | Production rate for $A\beta$ |
| $\text{clear}_{A\beta}$ | 3 | $\text{years}^{-1}$ | Clearance rate for $A\beta$ |
| $\text{trans}_{A\beta}$ | 3 | $\text{M}^{-1} \cdot \text{years}^{-1}$ | Transformation of $A\beta$ into its toxic isoform |
| $\text{clear}_{A\beta t}$ | 2.4 | $\text{years}^{-1}$ | Clearance rate for toxic $A\beta$ |
| $\text{prod}_T$ | 3 | $\text{M} \cdot \text{years}^{-1}$ | Production rate for tau |
| $\text{clear}_T$ | 3 | $\text{years}^{-1}$ | Clearance rate for tau |
| $\text{trans}_T$ | 3 | $\text{M}^{-1} \cdot \text{years}^{-1}$ | Transformation of tau into its toxic isoform |
| $\text{clear}_{Tt}$ | 2.55 | $\text{years}^{-1}$ | Clearance rate for toxic tau |
| $\text{syn}$ | 0.4 | $\text{M}^{-2} \cdot \text{years}$ | Synergistic effect between toxic $A\beta$ and toxic tau |
| $\rho$ | 50 | $\text{cm} \cdot \text{years}^{-1}$ | Effective diffusion constant |
| $A\beta_{\text{damrate}}$ | 1 | $\text{M}^{-1} \cdot \text{year}^{-1}$ | Damage rate for $A\beta t$ |
| $\text{TAU}_{\text{damrate}}$ | 1 | $\text{M}^{-1} \cdot \text{year}^{-1}$ | Damage rate for hyperphosphorilated TAUt |
| $\text{ha}_{\text{damrate}}$ | 5 | | Damage rate for hyperactivity |
| $cA\beta_{\text{exc}}$ | 0.8 | $\text{years}^{-1}$ | Constant for the effect of $A\beta t$ on excitation locally |
| $cA\beta_{\text{inh}}$ | 0.4 | $\text{years}^{-1}$ | Constant for the effect of $A\beta t$ on inhibition locally |
| $cT_{\text{exc}}$ | 1.8 | $\text{years}^{-1}$ | Constant for the effect of TAUt on excitatory local connectivity |
| $cT_{\text{inh}}$ | 1.8 | $\text{years}^{-1}$ | Constant for the effect of TAUt on inhibitory local connectivity |
| $cT_{\text{sc}}$ | 0.05 | $\text{years}^{-1}$ | Constant for the effect of TAUt on interregional connectivity |
| $H_{\text{e}_{\text{max}}}$ | 3.65 | | Maximum value allowed for $H_e$ parameter change |
| $C_{\text{ip}_{\text{min}}}$ | 13.25 | | Minimum value allowed for $C_{\text{ip}}$ parameter change |
| $C_{\text{ep}_{\text{min}}}$ | 28 | | Minimum value allowed for $C_{\text{ep}}$ parameter change |
| $T_{\text{sc}_{\text{max}}}$ | 0.3 | | Maximum damage of TAUt on interregional connectivity |

**Table 3.** Regions of the cingulum bundle included in the BNM Bubb et al. (2018). *Aβ* and tau superscripts denote the regions used as seeding (Palmqvist et al., 2017; Braak and Braak, 1991). I, II, III, and IV superscripts denote the regions included in the Braak stages (Therriault et al., 2022).

|  |  |
| --- | --- |
| Left Frontal Pole | Right Frontal Pole |
| Left Rostral Middle Frontal | Right Rostral Middle Frontal |
| Left Caudal Middle Frontal | Right Caudal Middle Frontal |
| Left Superior Frontal | Right Superior Frontal |
| Left Lateral Orbitofrontal $A\beta$ | Right Lateral Orbitofrontal $A\beta$ |
| Left Medial Orbitofrontal $A\beta$ | Right Medial Orbitofrontal $A\beta$ |
| Left Insula $A\beta$; IV | Right Insula $A\beta$; IV |
| Left Caudal Anterior Cingulate | Right Caudal Anterior Cingulate |
| Left Rostral Anterior Cingulate | Right Rostral Anterior Cingulate |
| Left Posterior Cingulate $A\beta$; IV | Right Posterior Cingulate $A\beta$; IV |
| Left Isthmus Cingulate $A\beta$ | Right Isthmus Cingulate $A\beta$ |
| Left Superior Parietal | Right Superior Parietal |
| Left Inferior Parietal $IV$ | Right Inferior Parietal |
| Left Precuneus $A\beta$ | Right Precuneus $A\beta$ |
| Left Inferior Temporal $IV$ | Right Inferior Temporal $IV$ |
| Left Parahippocampal $III$ | Right Parahippocampal $III$ |
| Left Hippocampus $II$ | Right Hippocampus $II$ |
| Left Thalamus | Right Thalamus |
| Left Amygdala $III$ | Right Amygdala $III$ |
| Left Entorhinal cortex $\tau$; I | Right Entorhinal cortex $\tau$; I |

Proteinopathy dynamics are described by the heterodimer model, one of the most common hypotheses that describe the prion-like spreading of toxic proteins. This hypothesis suggests that a healthy (properly folded) protein misfolds when it interacts with a toxic version of itself (misfolded; the prion/seed) following the latter’s structure as a template (Garzón et al., 2021). Therefore, this model includes healthy and toxic versions of amyloid-beta (*Aβ* / *Aβt*) and tau (TAU/TAUt) that are produced, cleared, trans-formed (from healthy to toxic), and propagated in the SC with N nodes.

The proteinopathy dynamics for i *∈* [1, N] were adapted from Alexandersen et al. (2023) to include the effect of hyperactivity on the enhanced production of *Aβ* (Cirrito et al., 2005; Yamamoto et al., 2015; Kamenetz et al., 2003; Busche and Hyman, 2020; Stargardt et al., 2015) and on the biased prion-like propagation of TAUt to hyperactive regions (Rodriguez et al., 2020); see Table 2 for a description of parameters):

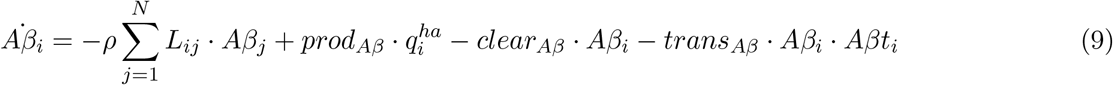

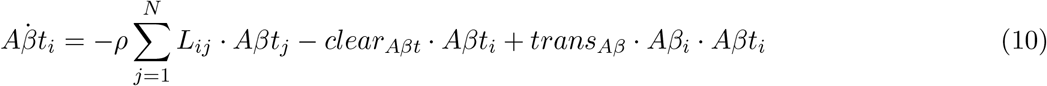

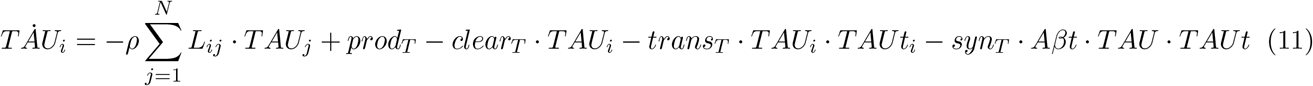

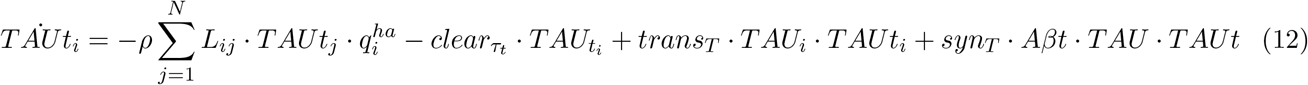

Where *prod, clear* and *trans* stand for rates of protein production, clearance, and transformation from a healthy isoform to a toxic one. *ρ* stands for a diffusion constant, and *L*_*ij*_ for the laplacian diffusion term:

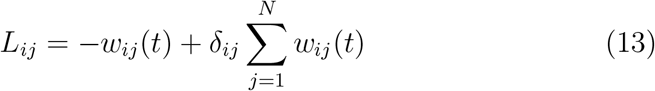

where *δ*_*ij*_ stands for the kronecher delta (i.e. the NxN identity matrix). The impact of protein concentration on the brain network model is established through two sets of transfer functions. First, a set of damage functions that are used to translate the concentration of toxic proteins into damage variables (Alexandersen et al., 2023):

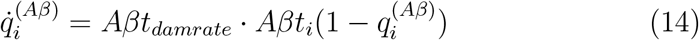

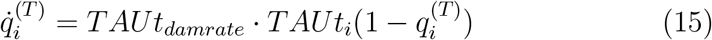

Second, a set of equations that translate the damage just mentioned into NMM parameter changes. We adapted these equations to the JR NMM. It was assumed that each protein (*Aβt* and TAUt) impacts the excitation/inhibition balance in different ways:

- *Aβt* generates neuronal hyperactivity through the impairment of GABAergic synapses (Garcia-Marin, 2009; Ulrich, 2015; Limon et al., 2012; Verret et al., 2012) and through the disruption of glutamate reuptake (Zott et al., 2019).
- Soluble hp-tau (TAUt) has been shown to change significantly the number and morphology of dendritic spines in pyramidal cells, producing neural silencing that dominates over the *Aβt* hyperactivation (Busche et al., 2019; Merino-Serrais et al., 2013).

The effects of *Aβt* will be modelled through changes in the local parameters of the JR NMM including an increase in H_e_ related to the disruption of glutamate reuptake (16), and a reduction of C_ip_ related to GABAergic deficits (eq. 17). The silencing effects of TAUt will be modelled through reductions in the local and interregional parameters that represent synaptic afferences to pyramidal cells, including C_ip_ and C_ep_ of local JR NMMs (eqs. 17 and 18) and the SC edge weights in the areas affected by TAUt (eqs. 19-20).

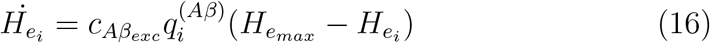

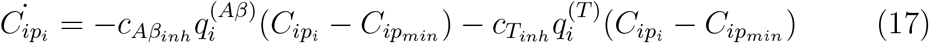

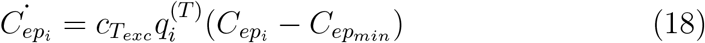

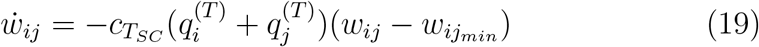

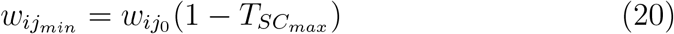

To this point, we have completed half of the closed-loop model. The second half regards the impact of the simulated neural activity on protein propagation. Neural hyperactivity fosters extracellular secretion and deposition of *Aβ* generating a cyclical phenomenon that reinforces bidirectionally neural hyperactivity and *Aβ* concentration (Cirrito et al., 2005; Yamamoto et al., 2015; Kamenetz et al., 2003; Busche and Hyman, 2020; Stargardt et al., 2015). Additionally, hyperactivity orients the prionic propagation of TAUt in the brain (Rodriguez et al., 2020) following the Braak stages (Braak and Braak, 1991).

To capture these effects in our model, we introduced an additional damage variable for hyperactivity (eq. 21). It was measured as the increase in firing rate from a baseline simulation in which proteinopathies do not affect (eq. 22). This damage variable was implemented in equations 9 and 12 by multiplying the production of *Aβ* and biasing the Laplacian term for TAUt distribution.

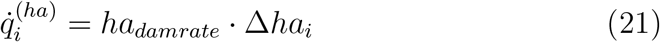

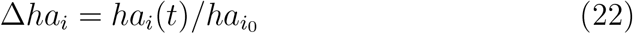

### 2.5 In-silico experiments and simulations

In this work, we performed simulations for several complementary in-silico experiments. First, we explored the parameter spaces of a single JR NMM to evaluate the effects of proteinopathy on neural activity, by varying the AD parameter candidates (H_e_, C_ip_, C_ep_, and w_ij_). As simulations were performed with a single node, we accounted for the reduction of interregional connectivity (w_ij_) produced by hp-tau using the mean input to the node (*p*). In total, we performed four sets of simulations in this in-silico experiment: the first one focused on the parameters related to *Aβ* effects, and the remaining three focused on parameters related to tau effects. The simulations were performed for 20 seconds of neural activity discarding the initial 12 to avoid transients, and we recorded information regarding the spectral frequency peak, spectral power in different frequency bands, and firing rate.

Second, we built and simulated the neurotoxicity model to deepen the mechanisms of AD progression. The initial conditions of the proteinopathy dynamics were fixed using a toxic protein seeding following literature on AD staging: the entorhinal cortex for tau (Braak and Braak, 1991) with an initial TAUt=0.0025 per region; and insula, precuneus, posterior and isthmus cingulate cortices, and orbitofrontal cortex for *Aβ* (Palmqvist et al., 2017) with an initial *Aβt*=0.0125. The proteinopathy dynamics were evaluated for each dt=0.25 (yrs) over 40 years, while the BNM was evaluated for each dt=1 (yrs) updating its parameters and simulating it for 10 seconds discarding the initial 2 of transients. Therefore, the transfer of information between the two parts of the model was executed with dt=1 (yrs).

We used the neurotoxicity model in several ways: first, we adjusted the model by exploring the impact of the limits of change for the parameter candidates 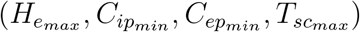 and two BNM free parameters (*g* and *s*) on the temporal evolution in terms of averaged frequency peak, relative alpha power, firing rate, and PLV calculated using the Phase Locking Value (PLV; (Lachaux et al., 1999; Bruña et al., 2018)) between signals filtered in the alpha band (8 -12 Hz):

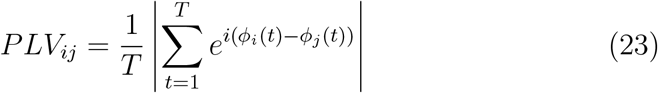

The adjusted parameters are included in Tables 1 and 2. Second, we simulated the model with the adjusted parameters to describe its outputs in terms of parameter trajectories (eqs. 16-19), spectra, firing rate, and PLV of the simulated brain regions; third, we disentangled the contribution of *Aβ* and hp-tau to the changes in inhibition isolating their respective effect in the model evolution using alternatively the parameters c*Aβ*_inh_and cT_inh_ equal to zero and computing the parameter space for the limit of change of C_ip_. And finally, we used it to evaluate the spatiotemporal evolution of the model with the Braak stages as a reference (see below).

### 2.6 Spatiotemporal profile assessment

We evaluated the spatial propagation of hp-tau using the Braak stages as a reference. Braak stages were adapted from (Therriault et al., 2022) considering the regions included in our cingulum bundle network (see Table 3): *stage one (rI)* included the entorhinal cortex; *stage two (rII)* included the hippocampus; *stage three (rIII)* included the parahippocampus and amygdala; *stage four (rIV)* included insula, posterior cingulate, inferior temporal, and inferior parietal; *stage five (rV)* included other cortical regions. Also, we evaluated the impact of *Aβ* seeding over the temporal antero-posterior differentiation in terms of firing rate and FC.

We simulated our closed-loop neurotoxicity model by implementing different seeding strategies. The baseline seeding (i.e., fixed strategy) was the same used in previous studies (Alexandersen et al., 2023; Thompson et al., 2020) as stated above. The remaining strategies consisted in randomizing the seeding of *Aβ*, randomizing the seeding of hp-tau, and randomizing both. The randomization respected two conditions: the number of seeded regions had to be the same as in the fixed strategy, and the seeding had to be symmetrical between hemispheres. For the antero-posterior differentiation, we also limited the randomization of *Aβ* seeding to posterior areas or anterior areas.

One hundred simulations were performed per seeding strategy. For each timestep, we averaged the concentration of hp-tau in the regions of the Braak stages to determine what Braak regions were presenting higher levels of the protein, and we sorted them into Braak sequences. Then, we extracted the dominant Braak sequence per simulation as the most repeated sequence in time. For the antero-posterior differentiation, we averaged posterior and anterior neural dynamics and we extracted the time to peak (given the rise and decay in the metrics) of each average.

### 2.7 Code Accessibility

The models described above were implemented in Python 3.9, and executed in a Surface Pro 7 (8Gb RAM) running Windows 11 and a HPC cluster. The code used in the paper is freely available online at [URL redacted for double-blind review]. The code is available as Extended Data.

## 3 Results

The proposed model for the physiological changes in AD is based on extensive literature that describes the effects of *Aβ* and hp-tau on neural tissue. These changes include the disruption of glutamate reuptake, the reduction of GABAergic synapses, and the reduction of pyramidal dendritic spines. To effectively model these changes, we proposed four parameter candidates that could account for the observed effects. Specifically, we selected two parameters to model the effects of *Aβ* on neural activity including the amplitude of the excitatory postsynaptic potential (H_e_) to account for the disruption of glutamate reuptake, and the number of local synaptic contacts from inhibitory interneurons to pyramidal neurons (C_ip_) to account for the reduction of GABAergic synapses. Regarding the effects of tau, we selected three parameter candidates that could effectively model the disruption of pyramidal dendritic spines: C_ip_ mentioned above, C_ep_ as the local synaptic contacts from excitatory interneurons to pyramidal neurons, and w_ij_ as the interregional SC weights.

### 3.1 Single-node experiments favour inhibition over excitation to explain hyperactivity

In order to gain a better understanding of the impact of toxic proteins on neural behaviour, we explored the parameter spaces of the proposed candidates (H_e_, C_ip_, C_ep_, and w_ij_) through single-node NMM simulations. Note that we used the mean input to the node (p) to account for the reduction of w_ij_ as simulations were performed with a single node.

In general terms, and in line with previous analysis of the JR model, results showed two main patterns of oscillation: a fast limit cycle in the alpha frequency band (8 -12 Hz), and a slow limit cycle in theta frequency band (4 -8 Hz). These two behaviours were categorically separated in the parameter spaces as can be observed in the frequency and relative power charts of Figure 1. The noisy regions surrounding the main limit cycles correspond to the fixed point states of JR that are found at both extremes of the excitation-inhibition continuum. In these states, the model might not have enough activation to give rise to an oscillatory behaviour, or it might be saturated by a too-high excitation. In both states, the noise acts as a driver giving rise to low-amplitude and low-frequency noisy oscillations, and the more stable the fixed point (i.e., more extreme excitation-inhibition balance) the lower the frequency and amplitude. Accordingly, note that the noisy regions coincide with the low-power regions in the absolute power plots of Figure 1.

**Figure 1.**
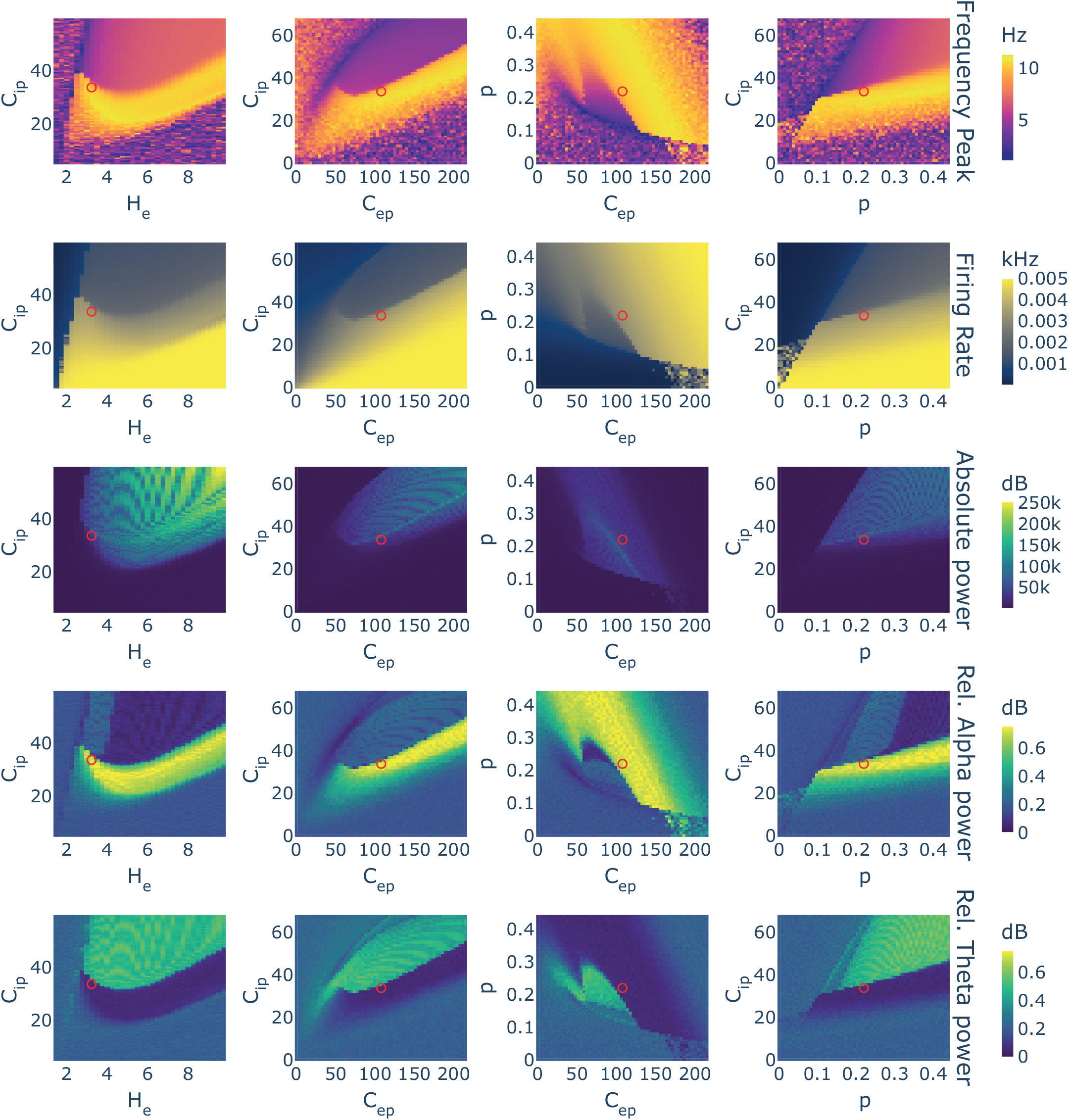
Single node in-silico experiment. The behaviour of a single neural mass model was simulated varying parameter values. Four different experiments are presented in columns exploring different combinations of parameters. In rows, measures derived from the simulations: oscillatory frequency peak of the node, mean firing rate, absolute power, and the relative power in two frequency bands: alpha and theta. The red circle points out the default parameters value. Simulations were performed without noise.

Focusing on the limit cycle regions, the parameter candidates related to intranode excitation (i.e., H_e_ and C_ep_) showed quadratic relationships in their parameter spaces, producing curved isopleth borders separating the slow and fast oscillatory behaviours (i.e., theta and alpha, respectively). These curves represented a convex relationship between the parameters and the neural behaviour in terms of frequency, power, and firing rate. The position of their original values derived from biology (Jansen and Rit, 1995) differed with respect to the vertex of the quadratic curve: the original H_e_ was found at lower values than the vertex, while C_ep_ was found at higher values.

For the other two parameters (i.e., C_ip_ and p) the observed changes inside the limit cycle regions were monotonical, and better described as logistic relationships in which the changes happen quickly at the edge between behaviours. These relationships were direct in the case of p with both frequency peak and firing rate, in contrast to the inverse relationship found for C_ip_ (i.e., lower inhibition raised the firing rate and the frequency of oscillation). Note that lowering inhibition out of the limit cycle regions had two different effects on firing rate and spectra. The firing rate kept rising monotonically as inhibition lowered, while the frequency peak and power tended to be reduced getting into the noisy low amplitude and slow region commented above.

In this model, two parameters could explain the hyperactivation produced by *Aβ*: C_ip_ lowering down due to the reduction of GABAergic synapses, and H_e_ rising due to the disruption of glutamate reuptake. Assuming that the original values of these parameters, which were based on biological observations (Jansen and Rit, 1995), are a good approximation of the underlying ground truth, our experiment would favour a hypothesis in which the hyperactivity found in AD is explained more likely by the reduction of C_ip_ than by the increase of H_e_ given that the latter would induce an initial reduction of activation levels that is not expected in AD (see the effects of lowering C_ip_ and rising H_e_ in Figure 1, second row, first column).

### 3.2 Closed-loop neurotoxicity model: from proteins to neural activity

To explore the link between proteinopathy and neural activity, we extended a previously published multiscale model of AD evolution (Alexandersen et al., 2023; Thompson et al., 2020) by implementing a biologically plausible NMM (Jansen and Rit, 1995) and establishing a bidirectional interaction between neural activity and proteinopathy. The proteinopathy starts from an initial distribution of toxic seeds, and during the temporal evolution of the model, the BNM integrates the effects of those proteins over neural tissue, which in turn influences the production of *Aβ* based on firing rate. The regional evolution of the proteinopathy follows a sigmoidal shape with a final slight decay in *Aβ* toxic concentration (see average curves in Figure 2A). The model captures the time delays between the rise of *Aβ* and hp-tau reported in the literature (Therriault et al., 2022). The effect of proteinopathy on neural activity was modelled through changes in the NMM parameters mediated by a damage variable (see Figure 2B). These changes were monotonical and followed the sigmoidal shape of the proteinopathy and its damage. The impact of *Aβ* over the neural tissue was modelled as a reduction of C_ip_ and an increase in H_e_, while the impact of hp-tau was modelled as reductions of the local and interregional connectivity parameters explored before (C_ip_, C_ep_, w_ij_; see Figure 2C). The changes in parameters due to the proteinopathy generated a rise and decay in firing rate over time (see Figure 2D) that affected the production of *Aβ* and the distribution of hp-tau through a hyperactivity damage variable (see Figure 2E).

**Figure 2.**
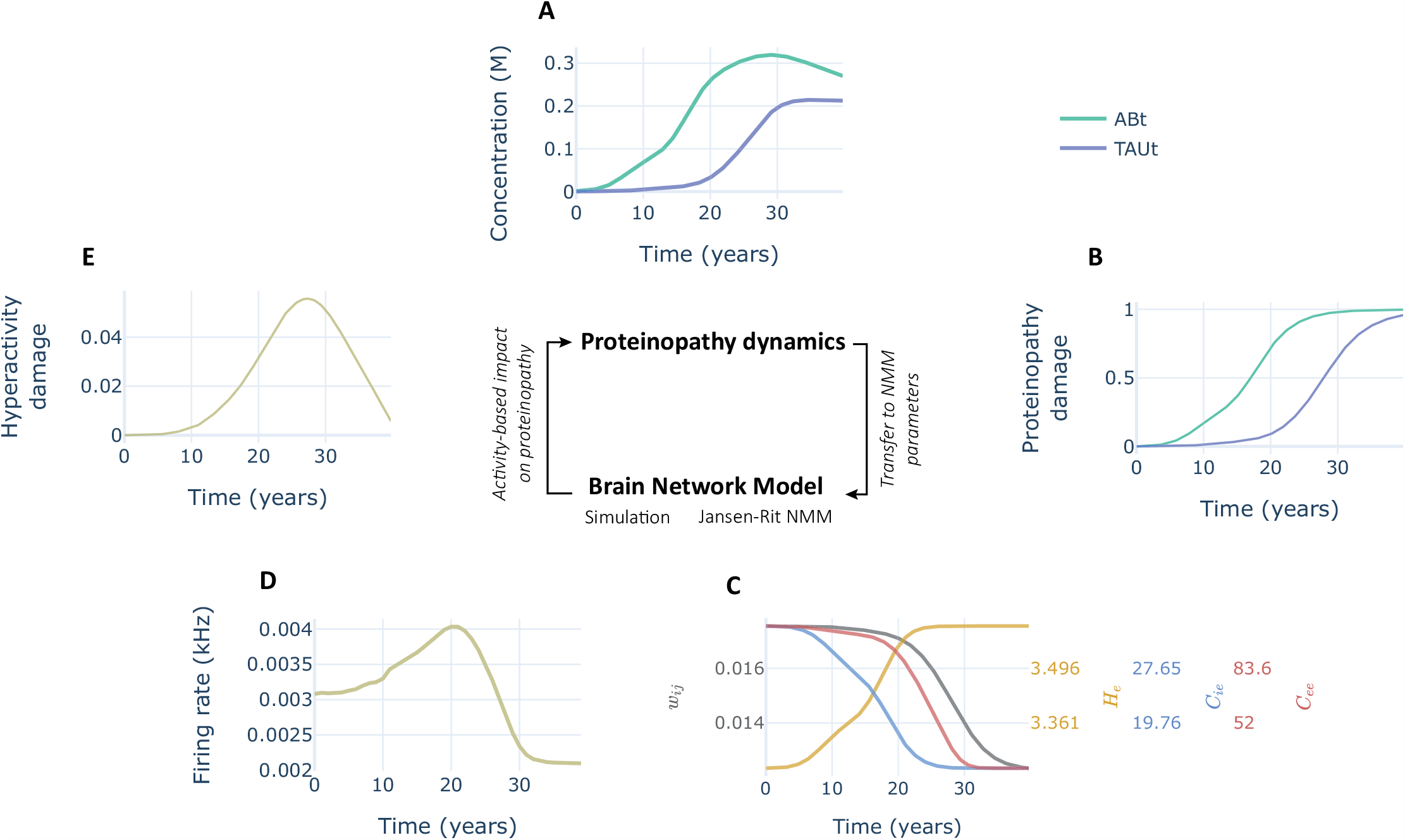
Parameters and state variables of the closed-loop neurotoxicity model. A) Averaged concentration of toxic proteins over time during model execution. B) Damage variables that determine the impact of the proteinopathy over the JR-BNM parameters. C) JR-BNM parameters change over time. D) Averaged firing rate derived from the simulation of the BNM. E) Damage variable that determines the impact of hyperactivity on the production and propagation of toxic proteins. The model is simulated following the proposed order.

The specific limits of change of the parameters (included in Table 2) were fitted by iterative explorations of the parameter spaces generated with the neurotoxicity model (e.g., Figure 4) to achieve the reproduction of some of the main pathophysiological changes of AD evolution (Maestú et al., 2021): frequency slowing, reduction in relative alpha power, hypo-/hyperactivity, and hypo-/hypersynchrony. Interestingly, we could not reproduce a clear temporal distinction between antero-posterior effects of the proteinopathy on FC (i.e., posterior regions are affected first) as it has been previously proposed (see Figure 3A-PLV). However, we also found slight antero-posterior differences in magnitude between those regions for FC and firing rate. Other neural activity outcomes of the model are shown in Figure 3 and will be carefully analyzed below along with the results of parameter space explorations.

**Figure 3.**
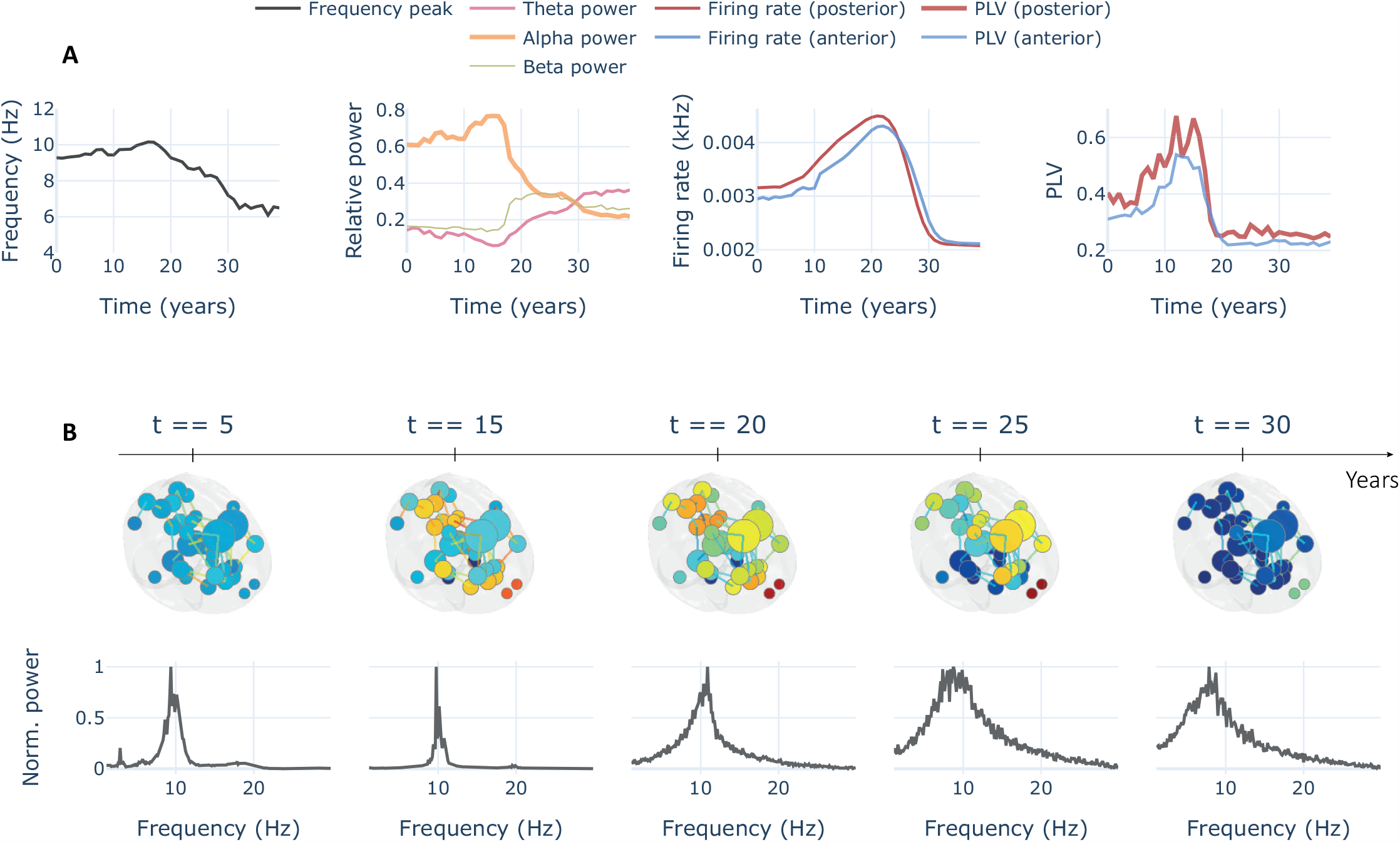
Functional measures from the closed-loop neurotoxicity model. A) Metrics extracted from the simulation of Fig. 2. We extracted spectral frequency peak, relative band power, firing rate, and FC from the simulation over 40 years of the neurotoxicity model. A quadratic relationship is found over time for all these metrics (i.e., measures increase early in time and decrease later). B) Temporal excerpts of the same simulation including 3D representations of the BNM with node size as degree, node colour as firing rate (orange/red is a higher rate), and edge colours as PLV. Last row shows averaged normalized spectra for all the regions included in the model.

**Figure 4.**
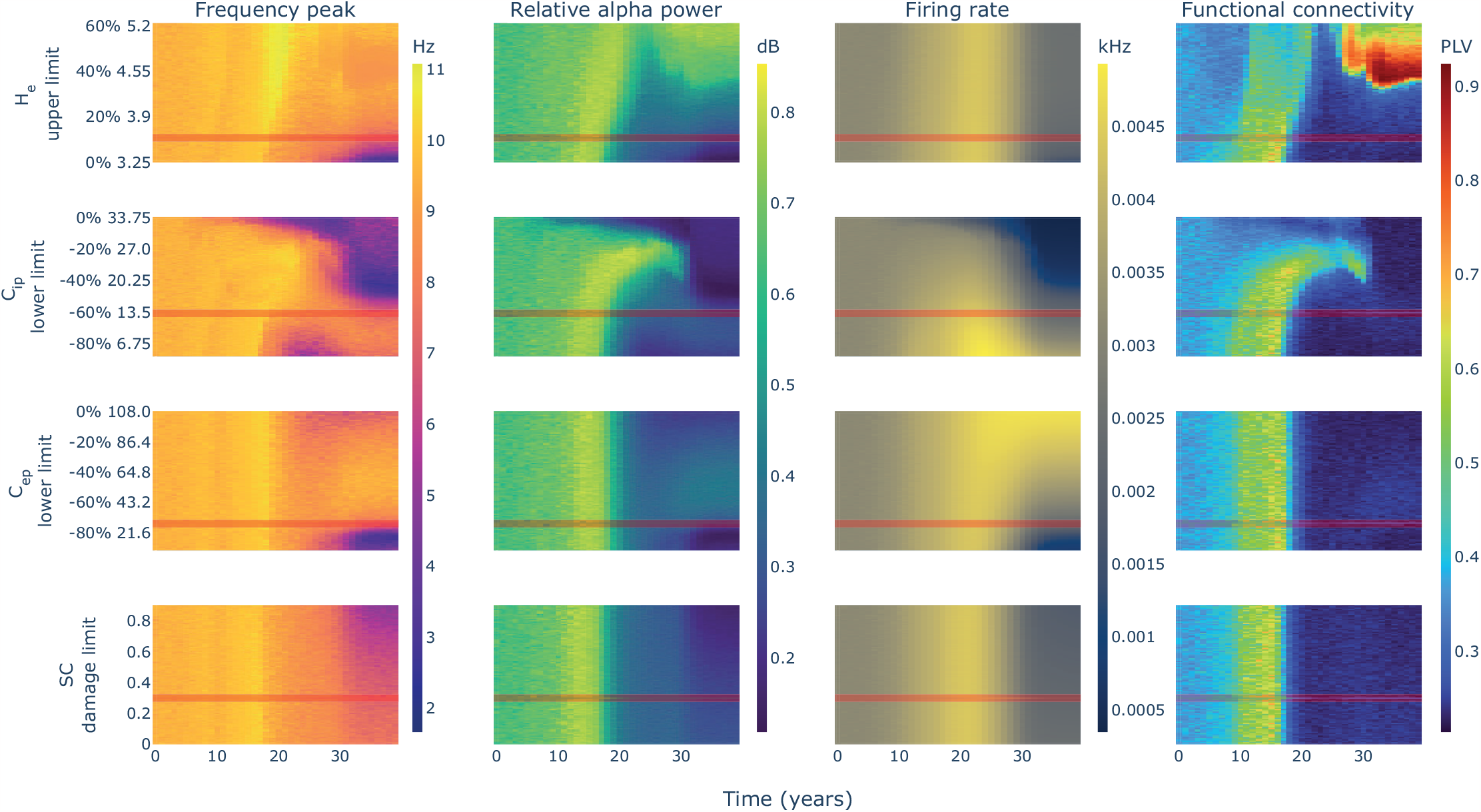
Parameter spaces for the limits of change of the JR-BNM parameter candidates. For each parameter candidate (rows) and limit value explored (heatmaps y-axis), one simulation of 40 years (heatmaps x-axis) is performed. For each simulation, four neural activity measures are extracted including averaged frequency peak, relative alpha power, firing rate, and functional connectivity (columns). When exploring a parameter candidate, all other parameters are kept fixed with a value included in Table 2 and highlighted in red in complementary spaces. Note how taking apart from zero the limits of change impact differently the resulting behaviour of the model. Two additional parameter spaces were computed to define values for g and s (see Figure 4-1).

### 3.3 Hyperactivity is not directly linked to hypersynchrony

The parameter space explorations that allowed us to fit the model are used in the following to establish relationships between different variables and outcomes derived from the neurotoxic model simulation. To obtain these parameter space explorations, we simulated the model with different values of the parameter candidates’ limits of change.

The model behaviour could be analyzed by dividing it into two stages: a first stage in which we find an increase in relative alpha power, firing rate, and FC; and a second stage in which all these measures decay (see Figure 4). These two stages are not synchronized for all measures. We observed a delayed peak in firing rate at the time interval of years 18 to 25, while all other measures peak earlier in time between years 10 and 18. From year 18 to 25, the firing rate keeps rising while power and FC have already lowered. This observation suggests a dissociation between firing rate and FC that has not been previously established.

### 3.4 The changes in inhibition lead the evolution of AD

Two parameter candidates could be responsible for the initial rising stage: the increase of H_e_ and the reduction of C_ip_. Looking at the parameter spaces, only the changes to the limits of C_ip_ showed a disruption in the rising phase when set below 20% of reduction (see Figure 4 second row). On the other hand, changes to the limits of H_e_ did not show any significant effect in this stage, even using an upper limit equal to 0% (i.e., keeping the parameter fixed in time) the rise in all measures could still be observed (see Figure 4, first row). This would suggest that the rising stage is mainly controlled in our model by the changes in inhibition (i.e., the reduction of C_ip_ parameter) representing the *Aβ* dependent reduction of GABAergic synapses.

The decaying stage required a more complex analysis. In the previously exposed single-node experiments, we observed that lowering inhibition showed a transition from the slow limit cycle, to the fast limit cycle, to the noisy low frequency and amplitude oscillation in which excitation-inhibition imbalance saturates the regional response and leads the JR NMMs to operate in a fixed point state (see Figure 1 and further explanations above). In that context, the reduction of C_ip_ itself could explain the decay stage for frequency peak and relative alpha power by entering the noisy region where lowering inhibition (i.e., further imbalance of excitation-inhibition) leads to a more stable JR fixed-point state and therefore to lower spectral frequencies. Interestingly, those spectral changes were time-locked with the changes in FC (see Figure 3 and Figure 4). Despite explaining the decay stage for those three measures, the reduction of C_ip_ could not explain the decay for firing rate. Lowering inhibition produced a monotonic ascent as could be expected from Figure 1 even in the noisy low amplitude region of the single-node parameter spaces.

The decay of firing rate could then be explained by another two parameter candidates related to the tau-derived reduction of pyramidal dendritic spines: the reduction of C_ep_, and the reduction of SC weights. The parameter spaces showed that only the reduction of local excitation (i.e., C_ep_) interacted with the firing rate (see Figure 4, third column). Disabling the change for C_ep_ (i.e., C_ep_ change limit of 0%) showed a plateau in firing rate during the decay stage, while activating its reduction showed a progressive effect on the decay of the firing rate (see Figure 4 third row, compare -40%, -60%, and -80% change limits). This effect was not found for different levels of damage to the SC weights.

### 3.5 Amyloid-beta and hp-tau could independently produce AD effects

From the analysis above, we can derive that the disruption of inhibition is a relevant driver of AD progress. Given that both *Aβ* and hp-tau contribute to the disruption of inhibition by reducing GABAergic synapses and reducing dendritic spines respectively, we wondered whether they contribute differently to the inhibitory-related effects reported above.

To disentangle their contribution, we computed an additional set of parameter spaces in which we evaluated independently the impact of the inhibitory disruption caused by each protein (Figure 5). We observed a clear similarity between the parameter space of *Aβ*-only effects and the previously shown in Figure 4 full model. The isolated effect of TAU could also generate a slight rise and decay in power, frequency and FC delayed from the *Aβ* effect and shorter in time, with a more moderate change in firing rate. This would suggest that both proteins could produce independently the effects observed in AD due to the disruption of inhibition, although with different timings and hyperactivity levels.

**Figure 5.**
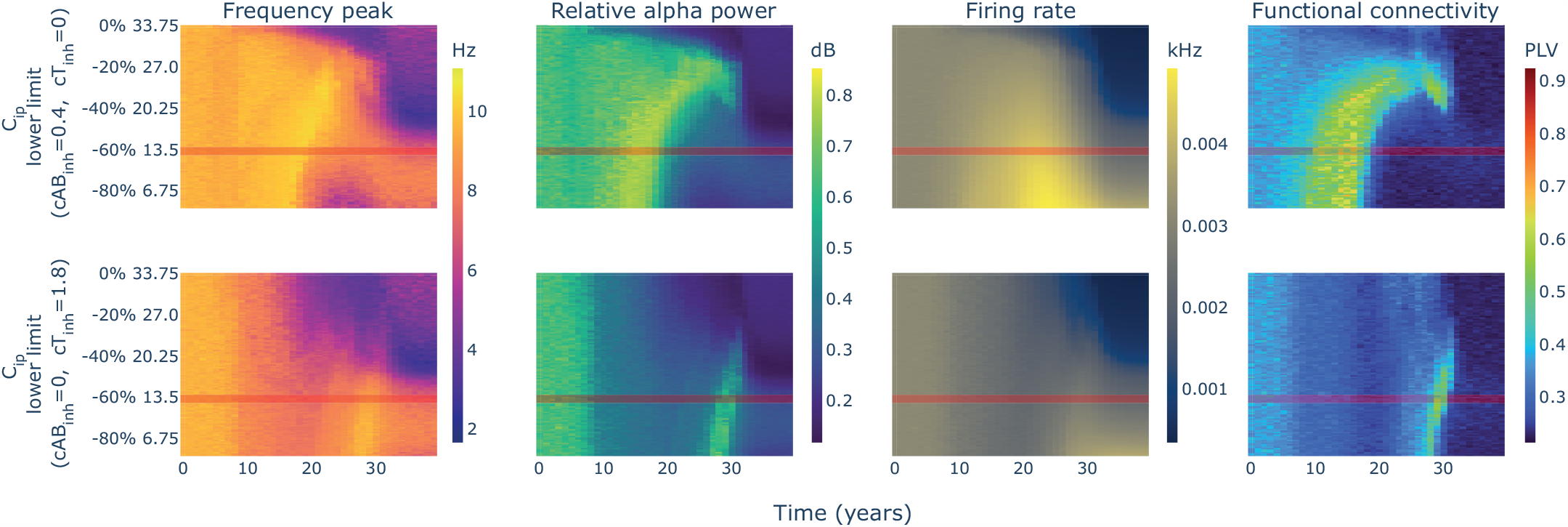
Parameter spaces for C_ip_ isolating the effects of *Aβ* and hp-tau. First row shows the results of simulating the neurotoxicity model with different limits of change for C_ip_ isolating the effects of *Aβ* on inhibition, while the second row shows the results of simulating the same parameter space isolating the effects of hp-tau on inhibition.

### 3.6 Toxic seeding determines the spatiotemporal profile of the evolution

The spatiotemporal sequence of hp-tau spreading in AD has been accurately characterized through the Braak stages. We used this characteristic spatiotemporal sequence to evaluate the spatial accuracy of the tauopathy simulated in our model, and the impact of the selected toxic proteins’ seeding regions on its spatial evolution. We implemented four seeding strategies including fixed seeding, *Aβ* randomized, hp-tau randomized and both randomized. We performed 100 simulations per strategy and we measured the concentration of hp-tau in the regions of the Braak stages extracting the dominant spatial sequence of propagation per simulation.

Results showed that fixed (i.e., as usual) and *Aβ* random seeding strategies reproduced effectively the Braak staging in 100% and 75% of the simulations, respectively (Figure 6A). In these simulations, just a slight temporal shift separated stages II (i.e., hippocampus) and III (i.e., parahippocampus and amygdala), and this way 25% of the simulations with *Aβ*-random seeding showed a sequence in which hp-tau reached stage III before stage II. Interestingly, randomizing tau seeding produced a variety of Braak sequences that lowered the dominance of the theoretical sequence to less than 10% of the simulations, at the same levels of randomizing both seeding locations. Examples of these sequences in time are shown in Figure 6B. These observations posit the importance of the localization of hp-tau seeds (i.e., early accumulations) to determine the evolution of its propagation.

**Figure 6.**
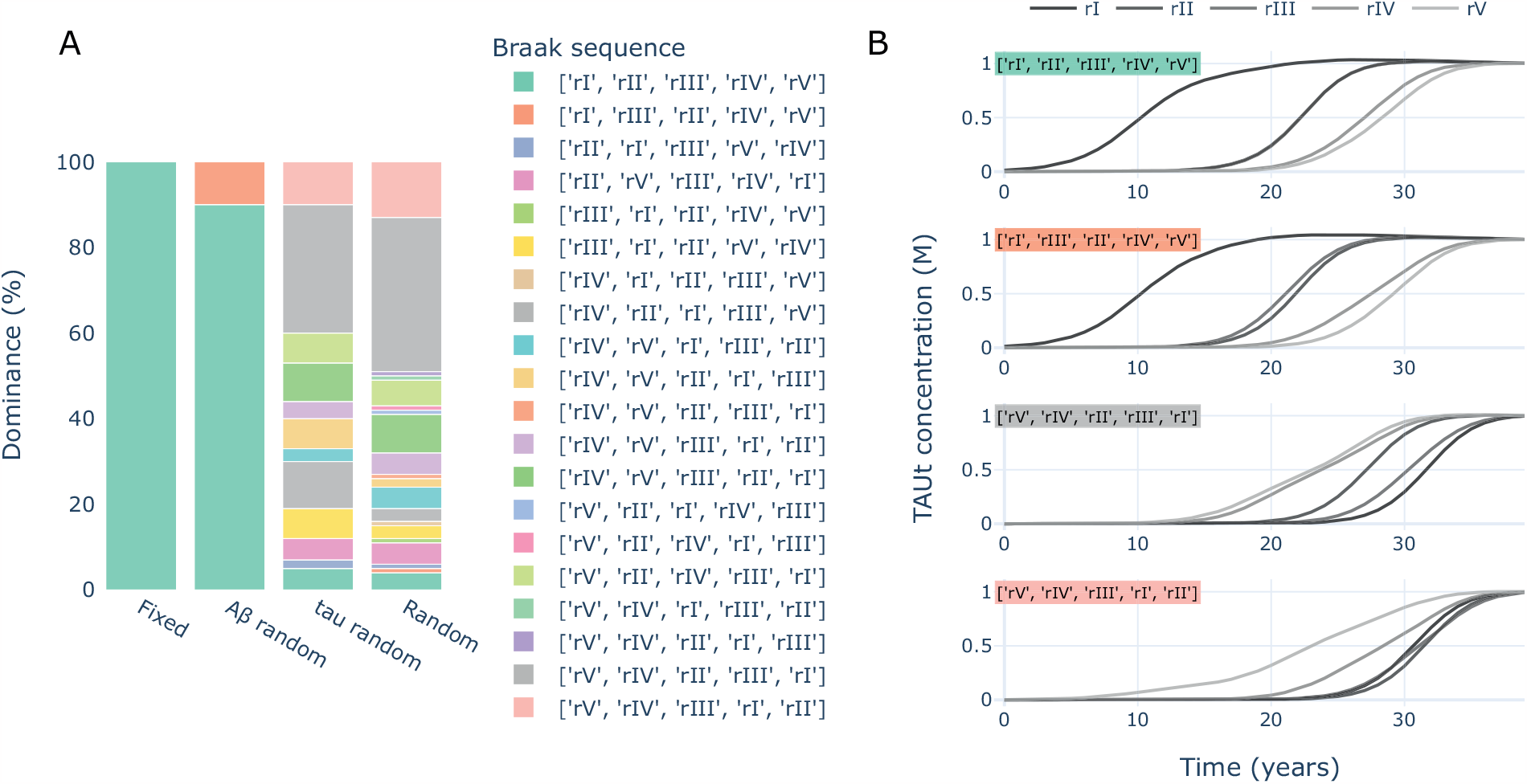
Dominance of Braak sequences per seeding strategy. A) The percentage of simulations per seeding mode in which different Braak sequences of hp-tau propagation were dominant. Fixed implied seeding a usual, *Aβ* random implied randomizing *Aβ* seeding and using the fixed version for hp-tau, vice versa for tau random, and Random implied randomizing both *Aβ* and hp-tau seeding. B) Samples of the most representative Braak sequences.

In addition, we wondered to what extent the antero-posterior temporal differentiation of the neurophysiological changes observed along the AD continuum could be related to the seeding of *Aβ*. We performed an additional set of simulations randomizing the seeding of *Aβ* and measuring the differences in firing rate and FC between anterior and posterior regions.

We observed a stable precession of posterior regions in terms of firing rate for the cases of fixed seeding (i.e., following Palmqvist et al. (2017)), for the randomized seeding in posterior regions, and for randomization over all regions (see Figure 7). Only in the case of anterior seeding, the model showed a similar timing between regions. This suggests that hyperactivity tends to appear earlier in posterior regions disregarding where *Aβ* starts to accumulate. Note that the seeding affected the levels of hyperactivity shown by the regions: non-seeded regions lowered their maximum firing rate (see Figure 7-1). In contrast, the antero-posterior temporal differentiation for FC relied more on seeding. When the seeding was located in posterior regions they peaked earlier while, on the contrary, when anterior regions were seeded they tended to peak earlier. Interestingly, these two strategies generated a general delay in the time to peak both in the seeded region and in the complementary.

**Figure 7.**
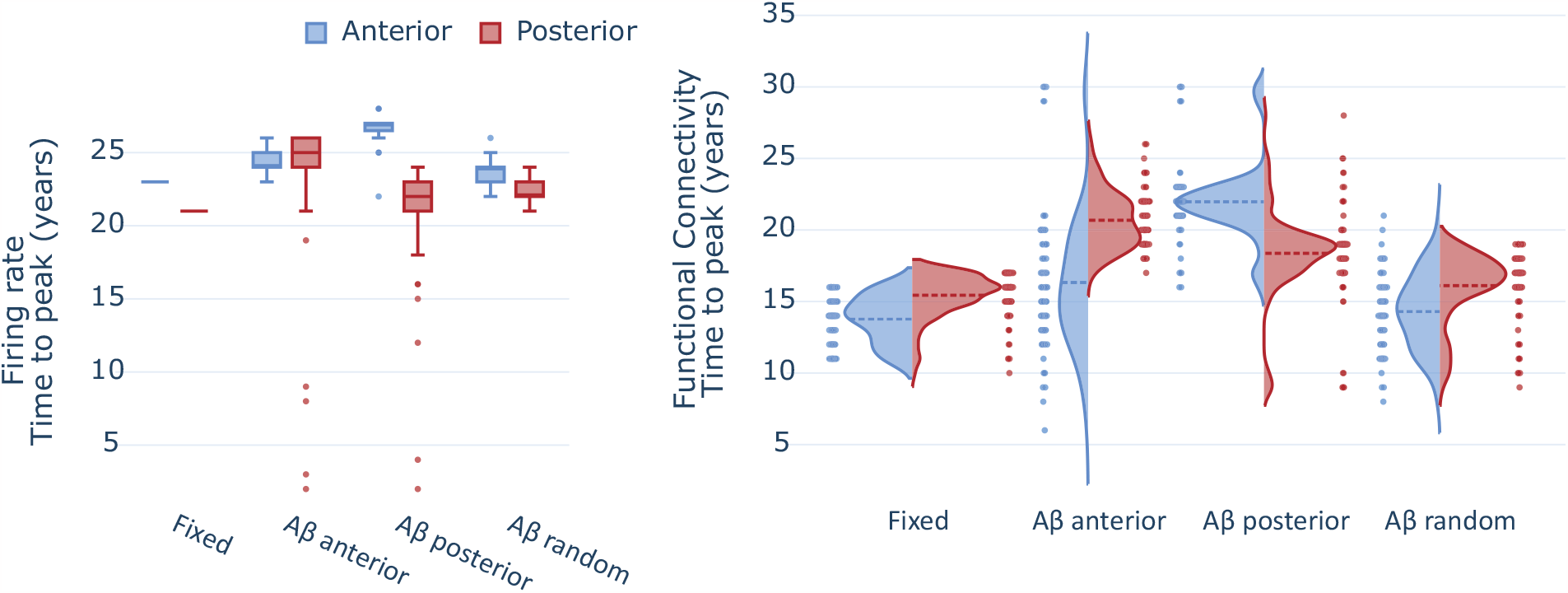
Impact of *Aβ* seeding on the antero-posterior temporal differentiation for firing rate (left) and FC (PLV; right). The neurotoxicity model shows a rise and decay for several variables including firing rate and FC. We measured the time elapsed to that peak for anterior and posterior regions as a proxy to understand where the neurophysiological changes happened before. Samples of the temporal evolution for each seeding strategy and for both firing rate and FC are represented in Figure 7-1.

## 4 Discussion

In this study, we developed a multiscale closed-loop neurotoxicity model of AD integrating a BNM that reproduced brain activity with another modelling the production and propagation of toxic proteins in the brain, both influencing each other through biologically plausible links. Our results showed that the disruption of inhibition was favoured to explain cellular hyperactivity, and indeed that inhibition was key to explaining the AD-related changes in frequency, power and FC. These changes in inhibition were primarily attributed to *Aβ*, although hp-tau alone could produce similar effects. Additionally, our results suggested a disconnection between cellular hyperactivity and interregional hypersynchrony. Finally, we observed a significant association between the spatiotemporal profile of the evolution of AD to the initial localization of the accumulation of AD proteins.

Single-node experiments favoured the lowering of inhibition (related to GABAergic synapses’ disruption) over the rising of excitation (related to impaired glutamate reuptake) to explain the characteristic hyperactivity produced by toxic *Aβ* on neural tissue. This idea is in line with previous animal studies showing a direct association between the levels of GABAergic synaptic activity and the hyperactivity found in the vicinity of amyloid plaques (Busche et al., 2008, 2015). Our result is based on the assumption that the biologically-informed original definition of the JR parameters is a good approximation to the characteristics of neural populations. Another definition of parameters could lead to different results. Nevertheless, it proposes a testable prediction regarding the extent to which these two *Aβ*-derived physiopathological phenomena may contribute to neural activation in AD.

Additionally, single-node experiments showed different effects for the changes implemented to the parameter candidates in terms of neural activity measures. Inside the limit cycle space, we observed quadratic relationships with firing rate and power for the local excitation parameters (i.e., H_e_ and C_ep_), while a monotonical relationship was found for the inhibitory parameter (i.e., C_ip_; Figure 1). These observations suggest two different roles for excitation and inhibition. While inhibition would predispose the system to order neural activation into synchronized oscillations, excitation is needed to trigger this mechanism. Therefore, both increasing and decreasing excitation levels could lead to a rise in cellular firing rate: while low excitation may become ineffective to trigger inhibition, high excitation could overcome the capacity of inhibition to organize action potentials and generate oscillatory behaviour. This type of contradictory effect (i.e., rising excitation parameter could lower firing rate, and vice versa) has been reported in empirical studies under the name of inhibitory stabilization of network dynamics, in which, for instance, the stimulation of inhibitory neurons produce a counterintuitive suppression of inhibitory firing (Sadeh and Clopath, 2020; Sanzeni et al., 2020). This mechanism is characteristic of networks with strong recurrent connectivity and protects the system from unstable dynamics such as epileptic seizures.

In agreement with single-node experiments, the simulation of the multiscale neurotoxicity model gave a prominent role to inhibition leading the neural activity changes typically observed in AD. Despite both *Aβ* and hp-tau can be associated with changes in inhibition -due to the disruption of GABAergic synapses, and to the reduction of dendritic spines that reduces the number of inhibitory inputs to pyramidal cells, respectively-, in the model, *Aβ* was linked to the main inhibitory changes that lead AD evolution. However, it was also shown that hp-tau alone could generate inhibitory effects in the same direction but weaker, shorter, and delayed in time.

The relationship between cellular hyperactivity and inter-regional hypersynchrony in AD has been suggested (Maestú et al., 2021; Sepulcre et al., 2017; Koelewijn et al., 2019) but not directly tested to our knowledge. The rationale behind this association is based on the studies linking *Aβ* independently with both hyperactivity and hypersynchrony. On one hand, some studies report higher cellular activation, or lower firing thresholds, in regions affected by *Aβ* (Palop and Mucke, 2010; Busche et al., 2008). On the other hand, some studies report spatial correlations between *Aβ* accumulation and increases of FC, which are mainly observed within the default mode network (Quevenco et al., 2020; Mormino et al., 2011; Schultz et al., 2017). From these observations, it has been derived that *Aβ* produces cellular hyperactivity, and that cellular hyperactivity, in turn, produces FC hypersynchrony. In our study, we show that hyper-activity and hypersynchrony might not be directly linked, as we found a reduction of FC while *Aβ* accumulation was still enhancing cellular hyper-activity, suggesting a temporal mismatch between the two. In contrast, FC levels were strongly associated in time with changes in spectral power. We believe that this dissociation between spectral power and firing rate is essential to understand the hyperactivity/hyperconnectivity question. Whenever hyperactivity overcomes the capacity of inhibition to order neural firing into an oscillatory behaviour, the regional activity may become noisier and therefore more difficult to synchronize interregionally. However, when hyperactivity moves the excitation/inhibition ratio towards a more balanced state, larger oscillations appear that may give rise to FC changes in a network.

The simulation of the closed-loop neurotoxicity model showed a sigmoidal shape in the evolution of the concentration of toxic proteins with a final slight decay for *Aβt* that is found in the empirical literature (Therriault et al., 2022). This decay is associated in our model with the reduction of hyperactivity that lowers the production of *Aβ* and therefore, leads to the recovery of the production-clearance balance. The impact of hyper-activity on the production of *Aβ* has been suggested in previous studies and linked to alterations in the endocytic machinery that could increase the rate of clathrin-mediated endocytosis of the amyloid precursor protein (APP), and also enlarge APP endosomes (Stargardt et al., 2015). After releasing these APP vesicles, a molecular cascade mediated by *β*-secretase and *γ*-secretase cleaves the molecule to generate *Aβ* (Haass and Selkoe, 2007).

An aspect that has not been captured by our model is the effect of chronic neuroinflammation. It may affect the normal functioning of glial cells, disrupting the clearance function of both microglia and astrocytes and, therefore, fostering protein accumulation and neuroinflammation (Henek et al., 2015; Henstridge et al., 2019). Although it is not the only additional aspect that could be implemented in the model due to its impact on AD progression (e.g., vascular damage leading to reduced protein clearance, infections such as herpes virus, genetic factors, etc.), we expect that by including these glial effects, we could reproduce the slight decay in hp-tau concentration that is found in literature, but not reproduced in our results. Further modelling studies including additional AD-related factors are needed.

Regarding simulated neural activity, our model qualitatively reproduced some of the most relevant biomarkers along the AD continuum, including the slowing of the alpha frequency, the rise and decay of relative alpha power, firing rate, and FC, and the Braak stages (Maestú et al., 2021). However, we could not directly reproduce the spatiotemporal dissociation in FC between the posterior and anterior brain regions that have been proposed in empirical research (Nakamura et al., 2017, 2018; Pusil et al., 2019; López-Sanz et al., 2016). This result was dependent on spatial seeding of *Aβ* that by including the orbitofrontal cortex, in addition to the precuneus and posterior cingulate cortex (Palmqvist et al., 2017), balanced anteroposterior changes in time. We believe that assuming this spatial seeding to be correct, the result could be expected empirically. However, we wonder whether the technical difficulties in detecting accumulations of *Aβ* could hinder the detection of earlier accumulations in posterior regions that could explain the electrophysiological evidence mentioned above.

This work provides a comprehensive and self-contained mechanistic explanation of how different factors contribute interactively to the course of AD and, therefore, it becomes relevant for urgent clinical needs. First, the observed hyperexcitation is the treatment target of the approved symptomatic drug Memantine (Reisberg et al., 2003, 2006) and other treatments currently under investigation such as the anti-epileptic drug Levetiracetam (Vossel et al., 2021). Several clinical trials in AD have determined that selecting the correct individuals at the correct disease stage for a particular intervention is crucial, our model aims to help in this process in the future. Further, both increased neural activity and altered synchrony can be targeted with invasive or non-invasive brain stimulation (Chang et al., 2018). Deep brain stimulation in the fornix has positively affected cognition, but mainly if the stimulation site precisely activates memory networks (Ríos et al., 2022). The promising effects of transcranial magnetic stimulation depend on a particular adaptation between the used frequencies and their effects on the excitation-inhibition balance (Weiler et al., 2019). Obviously, neuromodulation cannot only improve but also deteriorate the function of a diseased brain, and the prediction of an individual stimulation regime requires a deep understanding of the underlying mechanisms. The same applies to pharmacological treatments: while the recent success of Lecanemab in slowing down cognitive decline is promising and raises hopes for the amyloid hypothesis after decades (van Dyck et al., 2023), the observed effects are minimal, and the long-term outcome is still unclear. One avenue towards more efficient use of potentially disease-modifying treatments is, again, the choice of the correct timing for the therapy. The closed-loop-model in this work can even explain several potential reasons that hinder an effective slowing of the neurodegenerative process: it explains how both *Aβ* and hp-tau can drive the vicious circle independent from each other, and it formalizes how the self-sustaining process evolves even without the presence of the initiating factor. Altogether, only a holistic understanding of AD will open the possibility to once even preventing the development of dementia, and computational neuroscience can powerfully contribute to this.

### Extended Data 1

Code developed in Python 3.9 to simulate and visualize the closed-loop neurotoxicity model.

### Figure 4-1

Parameter spaces to select working point. Selected parameters were g=25 and s=20 m/s (see red highlights). Note how the rising of g leads to a situation in which no FC rise is observed, similar to the reduction of s, due to high early FC levels. Also, lowering g leads to the prebifurcation regime of the JR NMMs, a situation in which the spectral frequency peak lowers towards the delta band.

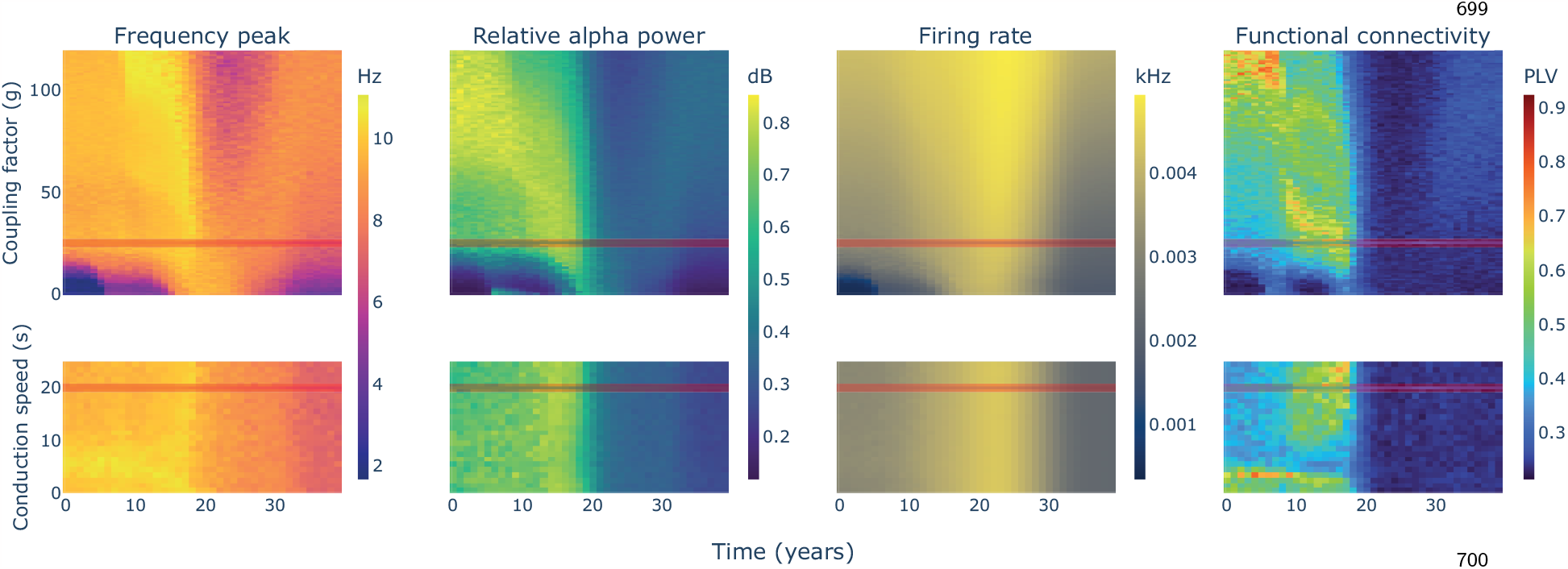

### Figure 7-1

Samples of the evolution for firing rate (left column) and FC (i.e., PLV; right column) in the antero-posterior differentiation experiments. In rows, each of the four seeding strategies implemented. The effect on FC is limited to a temporal shift of the curves, however, the seeding affects the level of hyperactivity reached by the anterior or posterior regions.

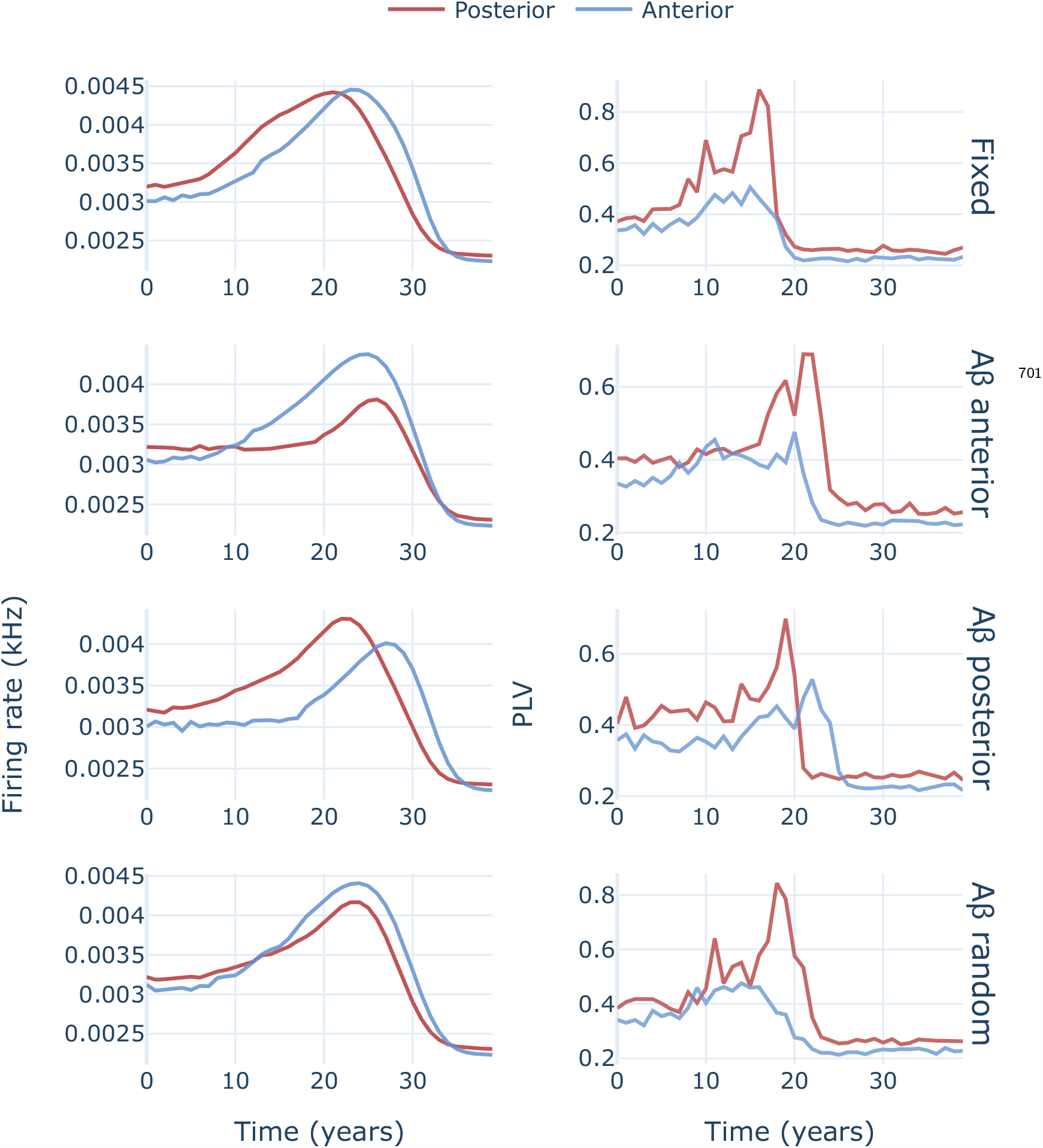

## References

Ahnaou, A., Moechars, D., Raeymaekers, L., Biermans, R., Manyakov, N. V., Bottelbergs, A., Wintmolders, C., Kolen, K. V., Casteele, T. V. D., Kemp, J. A., and Drinkenburg, W. H. (2017). Emergence of early alterations in network oscillations and functional connectivity in a tau seeding mouse model of alzheimer’s disease pathology. Scientific Reports, 7(1).

Alexandersen, C. G., de Haan, W., Bick, C., and Goriely, A. (2023). A multi-scale model explains oscillatory slowing and neuronal hyperactivity in Alzheimer’s disease. Journal of The Royal Society Interface, 20(198).

Braak, H. and Braak, E. (1991). Neuropathological stageing of Alzheimer-related changes. Acta Neuropathol, 82:239–59.

Braak, H., Thal, D. R., Ghebremedhin, E., and Tredici, K. D. (2011). Stages of the Pathologic Process in Alzheimer Disease: Age Categories From 1 to 100 Years. Journal of Neuropathology & Experimental Neurology, 70(11):960–969.

Breijyeh, Z. and Karaman, R. (2020). Comprehensive Review on Alzheimer’s Disease: Causes and Treatment. Molecules, 25(24):5789.

Bruña, R., Maestú, F., and Pereda, E. (2018). Phase locking value revisited: teaching new tricks to an old dog. Journal of Neural Engineering, 15(5):056011.

Bubb, E. J., Metzler-Baddeley, C., and Aggleton, J. P. (2018). The cingulum bundle: Anatomy function, and dysfunction. Neuroscience & Biobehavioral Reviews, 92:104–127.

Busche, M., Wegmann, S., Dujardin, S., Commins, C., Schiantarelli, J., Klickstein, N., Kamath, T., Carlson, G., Nelken, I., and Hyman, B. (2019). Tau impairs neural circuits, dominating amyloid-β effects, in Alzheimer models in vivo. Nat Neurosci, 22:57–64.

Busche, M. A., Eichhoff, G., Adelsberger, H., Abramowski, D., Wiederhold, K.-H., Haass, C., Staufenbiel, M., Konnerth, A., and Garaschuk, O. (2008). Clusters of hyperactive neurons near amyloid plaques in a mouse model of alzheimer’s disease. Science, 321(5896):1686–1689.

Busche, M. A. and Hyman, B. T. (2020). Synergy between amyloid-β and tau in Alzheimer’s disease. Nature Neuroscience, 23(10):1183–1193.

Busche, M. A., Kekuš, M., Adelsberger, H., Noda, T., Förstl, H., Nelken, I., and Konnerth, A. (2015). Rescue of long-range circuit dysfunction in alzheimer’s disease models. Nature Neuroscience, 18(11):1623–1630.

Chang, C.-H., Lane, H.-Y., and Lin, C.-H. (2018). Brain stimulation in alzheimer’s disease. Frontiers in Psychiatry, 9.

Chen, Y., Wang, Y., Song, Z., Fan, Y., Gao, T., and Tang, X. (2023). Abnormal white matter changes in Alzheimer’s disease based on diffusion tensor imaging: A systematic review. Ageing Res Rev, 87:101911.

Cirrito, J. R., Yamada, K. A., Finn, M. B., Sloviter, R. S., Bales, K. R., May, P. C., Schoepp, D. D., Paul, S. M., Mennerick, S., and Holtzman, D. M. (2005). Synaptic Activity Regulates Interstitial Fluid Amyloid-β Levels In Vivo. Neuron, 48(6):913–922.

Desikan, R., Ségonne, F., Fischl, B., Quinn, B., Dickerson, B., Blacker, D., Buckner, R., Dale, A., Maguire, R., Hyman, B., Albert, M., and Killiany, R. (2006). An automated labeling system for subdividing the human cerebral cortex on MRI scans into gyral based regions of interest. Neuroimage, 31:968–80.

Fischl, B., Salat, D., Busa, E., Albert, M., Dieterich, M., Haselgrove, C., van, d. K. A., Killiany, R., Kennedy, D., Klaveness, S., Montillo, A., Makris, N., Rosen, B., and Dale, A. (2002). Whole brain segmentation: automated labeling of neuroanatomical structures in the human brain. Neuron, 33:341–55.

Frost, B. and Diamond, M. I. (2009). Prion-like mechanisms in neurodegenerative diseases. Nature Reviews Neuroscience, 11(3):155–159.

Garcia-Marin, V. (2009). Diminished perisomatic GABAergic terminals on cortical neurons adjacent to amyloid plaques. Frontiers in Neuroanatomy, 3.

Garzón, D. N., Castillo, Y., Navas-Zuloaga, M. G., Darwin, L., Hardin, A., Culik, N., Yang, A., Castillo-Garsow, C., Ríos-Soto, K., Arriola, L., and Ghosh, A. (2021). Dynamics of prion proliferation under combined treatment of pharmacological chaperones and interferons. Journal of Theoretical Biology, 527:110797.

Gomez-Gutierrez, R. and Morales, R. (2020). The prion-like phenomenon in Alzheimer’s disease: Evidence of pathology transmission in humans. PLOS Pathogens, 16(10):e1009004.

Haass, C. and Selkoe, D. (2007). Soluble protein oligomers in neurodegeneration: lessons from the Alzheimer’s amyloid beta-peptide. Nat Rev Mol Cell Biol, 8:101–12.

Heneka, M. T., Carson, M. J., Khoury, J. E., Landreth, G. E., Brosseron, F., Feinstein, D. L., Jacobs, A. H., Wyss-Coray, T., Vitorica, J., Ransohoff, R. M., Herrup, K., Frautschy, S. A., Finsen, B., Brown, G. C., Verkhratsky, A., Yamanaka, K., Koistinaho, J., Latz, E., Halle, A., Petzold, G. C., Town, T., Morgan, D., Shinohara, M. L., Perry, V. H., Holmes, C., Bazan, N. G., Brooks, D. J., Hunot, S., Joseph, B., Deigendesch, N., Garaschuk, O., Boddeke, E., Dinarello, C. A., Breitner, J. C., Cole, G. M., Golenbock, D. T., and Kummer, M. P. (2015). Neuroinflammation in Alzheimer’s disease. The Lancet Neurology, 14(4):388–405.

Henstridge, C., Hyman, B., and Spires-Jones, T. (2019). Beyond the neuron-cellular interactions early in Alzheimer disease pathogenesis. Nat Rev Neurosci, 20:94–108.

Jansen, B. H. and Rit, V. G. (1995). Electroencephalogram and visual evoked potential generation in a mathematical model of coupled cortical columns. Biological Cybernetics, 73(4):357–366.

Kamenetz, F., Tomita, T., Hsieh, H., Seabrook, G., Borchelt, D., Iwatsubo, T., Sisodia, S., and Malinow, R. (2003). APP Processing and Synaptic Function. Neuron, 37(6):925–937.

Koelewijn, L., Lancaster, T. M., Linden, D., Dima, D. C., Routley, B. C., Magazzini, L., Barawi, K., Brindley, L., Adams, R., Tansey, K. E., Bompas, A., Tales, A., Bayer, A., and Singh, K. (2019). Oscillatory hyperactivity and hyperconnectivity in young APOE-ε4 carriers and hypoconnectivity in alzheimer’s disease. eLife, 8.

Lachaux, J., Rodriguez, E., Martinerie, J., and Varela, F. (1999). Measuring phase synchrony in brain signals. Hum Brain Mapp, 8:194–208.

Limon, A., Reyes-Ruiz, J. M., and Miledi, R. (2012). Loss of functional gaba(a) receptors in the alzheimer diseased brain. Proceedings of the National Academy of Sciences, 109(25):10071–10076.

López-Sanz, D., Bruña, R., Garcés, P., Camara, C., Serrano, N., Rodríguez-Rojo, I. C., Delgado, M. L., Montenegro, M., López-Higes, R., Yus, M., and Maestú, F. (2016). Alpha band disruption in the AD-continuum starts in the Subjective Cognitive Decline stage: a MEG study. Scientific Reports, 6(1).

Maestú, F., de Haan, W., Busche, M. A., and DeFelipe, J. (2021). Neuronal excitation/inhibition imbalance: core element of a translational perspective on Alzheimer pathophysiology. Ageing Research Reviews, 69:101372.

Masters, C. L., Bateman, R., Blennow, K., Rowe, C. C., Sperling, R. A., and Cummings, J. L. (2015). Alzheimer’s disease. Nature Reviews Disease Primers, 1(1).

Merino-Serrais, P., Benavides-Piccione, R., Blazquez-Llorca, L., Kastanauskaite, A., Rábano, A., Avila, J., and DeFelipe, J. (2013). The influence of phospho-tau on dendritic spines of cortical pyramidal neurons in patients with Alzheimer’s disease. Brain, 136(6):1913–1928.

Mormino, E. C., Smiljic, A., Hayenga, A. O., Onami, S. H., Greicius, M. D., Rabinovici, G. D., Janabi, M., Baker, S. L., Yen, I. V., Madison, C. M., Miller, B. L., and Jagust, W. J. (2011). Relationships between beta-amyloid and functional connectivity in different components of the default mode network in aging. Cerebral Cortex, 21(10):2399–2407.

Nakamura, A., Cuesta, P., Fernández, A., Arahata, Y., Iwata, K., Kuratsubo, I., Bundo, M., Hattori, H., Sakurai, T., Fukuda, K., Washimi, Y., Endo, H., Takeda, A., Diers, K., Bajo, R., Maestú, F., Ito, K., and Kato, T. (2018). Electromagnetic signatures of the preclinical and prodromal stages of Alzheimer’s disease. Brain, 141(5):1470–1485.

Nakamura, A., Cuesta, P., Kato, T., Arahata, Y., Iwata, K., Yamagishi, M., Kuratsubo, I., Kato, K., Bundo, M., Diers, K., Fernández, A., Maestú, F., and Ito, K. (2017). Early functional network alterations in asymptomatic elders at risk for Alzheimer’s disease. Scientific Reports, 7(1).

Palmqvist, S., Schöll, M., Strandberg, O., Mattsson, N., Stomrud, E., Zetterberg, H., Blennow, K., Landau, S., Jagust, W., and Hansson, O. (2017). Earliest accumulation of β-amyloid occurs within the default-mode network and concurrently affects brain connectivity. Nature Communications, 8(1).

Palop, J. J., Chin, J., and Mucke, L. (2006). A network dysfunction perspective on neurodegenerative diseases. Nature, 443(7113):768–773.

Palop, J. J. and Mucke, L. (2010). Amyloid-β induced neuronal dysfunction in alzheimer’s disease: from synapses toward neural networks. Nature Neuroscience, 13(7):812–818.

Pereira, J. B., Ossenkoppele, R., Palmqvist, S., Strandberg, T. O., Smith, R., Westman, E., and Hansson, O. (2019). Amyloid and tau accumulate across distinct spatial networks and are differentially associated with brain connectivity. eLife, 8.

Pusil, S., López, M. E., Cuesta, P., Bruña, R., Pereda, E., and Maestú, F. (2019). Hypersynchronization in mild cognitive impairment: the ‘X’ model. Brain, 142(12):3936–3950.

Quevenco, F. C., van Bergen, J. M., Treyer, V., Studer, S. T., Kagerer, S. M., Meyer, R., Gietl, A. F., Kaufmann, P. A., Nitsch, R. M., Hock, C., and Unschuld, P. G. (2020). Functional brain network connectivity patterns associated with normal cognition at old-age, local β-amyloid, tau, and APOE4. Frontiers in Aging Neuroscience, 12.

Ranasinghe, K. G., Hinkley, L. B., Beagle, A. J., Mizuiri, D., Dowling, A. F., Honma, S. M., Finucane, M. M., Scherling, C., Miller, B. L., Nagarajan, S. S., and Vossel, K. A. (2014). Regional functional connectivity predicts distinct cognitive impairments in alzheimer’s disease spectrum. NeuroImage: Clinical, 5:385–395.

Ranasinghe, K. G., Verma, P., Cai, C., Xie, X., Kudo, K., Gao, X., Lerner, H., Mizuiri, D., Strom, A., Iaccarino, L., Joie, R. L., Miller, B. L., Gorno-Tempini, M. L., Rankin, K. P., Jagust, W. J., Vossel, K., Rabinovici, G. D., Raj, A., and Nagarajan, S. S. (2022). Altered excitatory and inhibitory neuronal subpopulation parameters are distinctly associated with tau and amyloid in alzheimer’s disease. eLife, 11.

Reisberg, B., Doody, R., Stöffler, A., Schmitt, F., Ferris, S., and Möbius, H. J. (2003). Memantine in moderate-to-severe alzheimer’s disease. New England Journal of Medicine, 348(14):1333–1341.

Reisberg, B., Doody, R., Stöffler, A., Schmitt, F., Ferris, S., and Möbius, H. J. (2006). A 24-week open-label extension study of memantine in moderate to severe alzheimer disease. Archives of Neurology, 63(1):49.

Ríos, A. S., Oxenford, S., Neudorfer, C., Butenko, K., Li, N., Rajamani, N., Boutet, A., Elias, G. J. B., Germann, J., Loh, A., Deeb, W., Wang, F., Setsompop, K., Salvato, B., de Almeida, L. B., Foote, K. D., Amaral, R., Rosenberg, P. B., Tang-Wai, D. F., Wolk, D. A., Burke, A. D., Salloway, S., Sabbagh, M. N., Chakravarty, M. M., Smith, G. S., Lyketsos, C. G., Okun, M. S., Anderson, W. S., Mari, Z., Ponce, F. A., Lozano, A. M., and Horn, A. (2022). Optimal deep brain stimulation sites and networks for stimulation of the fornix in alzheimer’s disease. Nature Communications, 13(1).

Rodriguez, G. A., Barrett, G. M., Duff, K. E., and Hussaini, S. A. (2020). Chemogenetic attenuation of neuronal activity in the entorhinal cortex reduces Aβ and tau pathology in the hippocampus. PLOS Biology, 18(8):e3000851.

Sadeh, S. and Clopath, C. (2020). Inhibitory stabilization and cortical computation. Nature Reviews Neuroscience, 22(1):21–37.

Sanz-Leon, P., Knock, S. A., Spiegler, A., and Jirsa, V. K. (2015). Mathematical framework for large-scale brain network modeling in The Virtual Brain. NeuroImage, 111:385–430.

Sanzeni, A., Akitake, B., Goldbach, H. C., Leedy, C. E., Brunel, N., and Histed, M. H. (2020). Inhibition stabilization is a widespread property of cortical networks. eLife, 9.

Schultz, A. P., Chhatwal, J. P., Hedden, T., Mormino, E. C., Hanseeuw, B. J., Sepulcre, J., Huijbers, W., LaPoint, M., Buckley, R. F., Johnson, K. A., and Sperling, R. A. (2017). Phases of hyperconnectivity and hypoconnectivity in the default mode and salience networks track with amyloid and tau in clinically normal individuals. The Journal of Neuroscience, 37(16):4323–4331.

Sepulcre, J., Sabuncu, M. R., Li, Q., Fakhri, G. E., Sperling, R., and Johnson, K. A. (2017). Tau and amyloid β proteins distinctively associate to functional network changes in the aging brain. Alzheimer’s & Dementia, 13(11):1261–1269.

Stargardt, A., Swaab, D. F., and Bossers, K. (2015). The storm before the quiet: neuronal hyperactivity and Aβ in the presymptomatic stages of Alzheimer’s disease. Neurobiology of Aging, 36(1):1–11.

Stefanovski, L., Meier, J. M., Pai, R. K., Triebkorn, P., Lett, T., Martin, L., Bülau, K., Hofmann-Apitius, M., Solodkin, A., McIntosh, A. R., and Ritter, P. (2021). Bridging Scales in Alzheimer’s Disease: Biological Framework for Brain Simulation With The Virtual Brain. Frontiers in Neuroinformatics, 15.

Stefanovski, L., Triebkorn, P., Spiegler, A., Diaz-Cortes, M.-A., Solodkin, A., Jirsa, V., McIntosh, A. R., and and, P. R. (2019). Linking Molecular Pathways and Large-Scale Computational Modeling to Assess Candidate Disease Mechanisms and Pharmacodynamics in Alzheimer’s Disease. Frontiers in Computational Neuroscience, 13.

Therriault, J., Pascoal, T. A., Lussier, F. Z., Tissot, C., Chamoun, M., Bezgin, G., Servaes, S., Benedet, A. L., Ashton, N. J., Karikari, T. K., Lantero-Rodriguez, J., Kunach, P., Wang, Y.-T., Fernandez-Arias, J., Massarweh, G., Vitali, P., Soucy, J.-P., Saha-Chaudhuri, P., Blennow, K., Zetterberg, H., Gauthier, S., and Rosa-Neto, P. (2022). Biomarker modeling of Alzheimer’s disease using PET-based Braak staging. Nature Aging, 2(6):526–535.

Thompson, T. B., Chaggar, P., Kuhl, E., and and, A. G. (2020). Protein-protein interactions in neurodegenerative diseases: A conspiracy theory. PLOS Computational Biology, 16(10):e1008267.

Ulrich, D. (2015). Amyloid-Impairs Synaptic Inhibition via GABAA Receptor Endocytosis. Journal of Neuroscience, 35(24):9205–9210.

van Dyck, C. H., Swanson, C. J., Aisen, P., Bateman, R. J., Chen, C., Gee, M., Kanekiyo, M., Li, D., Reyderman, L., Cohen, S., Froelich, L., Katayama, S., Sabbagh, M., Vellas, B., Watson, D., Dhadda, S., Irizarry, M., Kramer, L. D., and Iwatsubo, T. (2023). Lecanemab in Early Alzheimer’s Disease. New England Journal of Medicine, 388(1):9–21.

van Nifterick, A. M., Gouw, A. A., van Kesteren, R. E., Scheltens, P., Stam, C. J., and de Haan, W. (2022). A multiscale brain network model links Alzheimer’s disease-mediated neuronal hyperactivity to large-scale oscillatory slowing. Alzheimer’s Research & Therapy, 14(1).

Verret, L., Mann, E. O., Hang, G. B., Barth, A. M., Cobos, I., Ho, K., Devidze, N., Masliah, E., Kreitzer, A. C., Mody, I., Mucke, L., and Palop, J. J. (2012). Inhibitory interneuron deficit links altered network activity and cognitive dysfunction in alzheimer model. Cell, 149(3):708–721.

Vossel, K., Ranasinghe, K. G., Beagle, A. J., La, A., Pook, K. A., Castro, M., Mizuiri, D., Honma, S. M., Venkateswaran, N., Koestler, M., Zhang, W., Mucke, L., Howell, M. J., Possin, K. L., Kramer, J. H., Boxer, A. L., Miller, B. L., Nagarajan, S. S., and Kirsch, H. E. (2021). Effect of levetiracetam on cognition in patients with alzheimer disease with and without epileptiform activity. JAMA Neurology, 78(11):1345.

Walker, L. C. (2018). Prion-like mechanisms in Alzheimer disease. In Human Prion Diseases, pages 303–319. Elsevier.

Weiler, M., Stieger, K. C., Long, J. M., and Rapp, P. R. (2019). Transcranial magnetic stimulation in alzheimer’s disease: Are we ready? eneuro, 7(1):ENEURO.0235–19.2019.

Yamamoto, K., ichi Tanei, Z., Hashimoto, T., Wakabayashi, T., Okuno, H., Naka, Y., Yizhar, O., Fenno, L. E., Fukayama, M., Bito, H., Cirrito, J. R., Holtzman, D. M., Deisseroth, K., and Iwatsubo, T. (2015). Chronic Optogenetic Activation Augments Aβ Pathology in a Mouse Model of Alzheimer Disease. Cell Reports, 11(6):859–865.

Yeh, F.-C. (2020). Shape analysis of the human association pathways. NeuroImage, 223:117329.

Yeh, F.-C., Liu, L., Hitchens, T. K., and Wu, Y. L. (2016). Mapping immune cell infiltration using restricted diffusion MRI. Magnetic Resonance in Medicine, 77(2):603–612.

Yeh, F.-C., Verstynen, T. D., Wang, Y., Fernández-Miranda, J. C., and Tseng, W.-Y. I. (2013). Deterministic Diffusion Fiber Tracking Improved by Quantitative Anisotropy. PLoS ONE, 8(11):e80713.

Yeh, F.-C., Wedeen, V. J., and Tseng, W.-Y. I. (2010). Generalized $ {q} $ -Sampling Imaging. IEEE Transactions on Medical Imaging, 29(9):1626–1635.

Yu, J., Lam, C., and Lee, T. (2017). White matter microstructural ab-normalities in amnestic mild cognitive impairment: A meta-analysis of whole-brain and ROI-based studies. Neurosci Biobehav Rev, 83:405–416.

Zott, B., Simon, M. M., Hong, W., Unger, F., Chen-Engerer, H.-J., Frosch, M. P., Sakmann, B., Walsh, D. M., and Konnerth, A. (2019). A vicious cycle of β amyloid–dependent neuronal hyperactivation. Science, 365(6453):559–565.

